# Targeting the mitochondrial RNA methyltransferase TRMT61B reveals new therapeutic opportunities in aneuploid cancer cells

**DOI:** 10.1101/2021.04.25.441348

**Authors:** Alberto Martín, Borja Vilaplana-Marti, Rocío IR Macías, Ángel Martínez-Ramírez, Ana Cerezo, Pablo Cabezas-Sainz, Maria Garranzo Asensio, Carolina Epifano, Sandra Amarilla, Déborah Gómez-Domínguez, Iván Hernández, Eduardo Caleiras, Jordi Camps, Rodrigo Barderas, Laura Sánchez, Susana Velasco, Ignacio Pérez de Castro

## Abstract

Chromosomal instability (CIN) is an important source of genetic and phenotypic variation that has been extensively reported as a critical cancer related property that improves tumor cell adaptation and survival. CIN and its immediate consequence, aneuploidy, provoke adverse effects on cellular homeostasis that need to be overcome by developing efficient anti-stress mechanisms. Perturbations in these safeguard responses might be detrimental for cancer cells and represent an important tumor specific Achilles heel since CIN and aneuploidy are very rare events in normal cells. On the other hand, epitranscriptomic marks catalyzed by different RNA modifying enzymes have been found to change under several stress insults. Although CIN and aneuploidy are important intracellular stressors, their biological connection with RNA modifications is pending to be determined. In an *in silico* search for new cancer biomarkers, we have identified TRMT61B, a mitochondrial RNA methyltransferase enzyme, to be associated with high levels of aneuploidy. In the present work, we study the connection of this molecule with cancer and aneuploidy. First, we show increased protein amounts of TRMT61B in tumor cell lines with imbalanced karyotype as well as in different tumor types compared to unaffected control tissues. In addition, we demonstrate that depletion of TRMT61B in melanoma cells reduces cell proliferation either by fostering apoptosis and inhibiting autophagy in high-aneuploid (ANE^high^) cells or by inducing senescence in the case of low-aneuploid (ANE^low^) cell lines. Further, TRMT61B elimination compromises mitochondrial function and reduces the expression of several mitochondrial encoded proteins that are part of the electron transport chain. Finally, transwell and xenograft experiments revealed a reduced invasive and tumorigenic capacity upon TRMT61B depletion that strengthen the therapeutic value of this aneuploidy-associated biomarker. These results, which connect tumorigenesis, aneuploidy and mitochondrial RNA methylation, bring to the cancer field a new putative strategy to specifically target high aneuploid tumors.

## Introduction

Chromosomal instability (CIN) is defined as a dynamic condition by which cells gain or lose whole or parts of chromosomes with each cell division. This ongoing propensity to alter the normal set of chromosomes is very common in the course of oncogenesis ^1,2^. In fact, aneuploidy, the immediate consequence of CIN that describes an imbalanced karyotype, is frequently detected in a high fraction of solid and hematopoietic tumors ^2,3^. Although not identical, CIN and aneuploidy are closely related events, so both terms will be used along the present work for introductory and discussion aspects. CIN might be considered as a pivotal hallmark of cancer disease that favors the acquisition of other cancer hallmarks required for tumorigenesis ^4^. Importantly, the genetic diversity associated with CIN might account for the development of drug resistance, high metastatic incidence and low survival rates of many cancer patients ^2,5–8^.

Paradoxically, in the short term, genomic disruption in general and karyotypic changes in particular, normally results in detrimental proliferative and metabolic effects due to different types of stresses, such as proteoxic and genotoxic burden, overloaded endoplasmic reticulum, augmented glycolytic flux and mitochondrial activity, and an increase of reactive oxygen species (ROS) ^9–14^. However, cancer cells are generally characterized by an aneuploid and proliferative phenotype which suggests that gain or loss of chromosomes can be finally advantageous for adaptation once the initial challenge of gene imbalance has been overcome. In fact, different studies showed that CIN can generates massive genetic and phenotypic variation that could drive rapid evolution to select cells that can tolerate high stress conditions and adapt to an ever-changing environment ^15–18^. To this regard, a plethora of cellular stress responses has been suggested to be adopted in response to intracellular and extracellular perturbations that might allow aneuploidy cells outcompete euploid counterparts in certain challenging environments and become dominant. These include, among others, upregulation of autophagy, DNA repair mechanisms, and antioxidant levels as well as more efficient bioenergetic and biosynthetic processes ^12,19–23^. In this sense, the gene dosage alterations and high ROS levels exhibited by CIN^+^ cells, impose an energy deficit related to additional protein synthesis, folding and clearance demands that can be satisfied with increased glycolytic and mitochondrial activities ^24–28^. Recent reports highlight that, in addition to a strong dependence on glycolysis, many tumors rely on oxidative phosphorylation for both ATP and macromolecule building block synthesis ^29^. However, this metabolic reprogramming is part of the solution but at the same time part of the problem, since an enhanced mitochondrial activity is one of the major sources of ROS production responsible for many deleterious effects on cell survival ^30–32^.

Therefore, safeguard mechanisms should be launched against redox imbalance and intracellular damage suggesting that different cellular stress responses need to work in a complementary fashion ^11,28^. Nevertheless, how aneuploidy can overcome deleterious effects and finally provide a selective advantage remains incompletely understood ^5^.

Over 160 RNA modifications involving post-transcriptional changes in the chemical composition of different classes of nuclear and mitochondrial RNAs have been described to date, with tRNAs and rRNAs containing the most numerous and chemically diverse modified ribonucleosides ^33,34^. These modifications range from the addition of a single methyl group in a nitrogenous base or sugar residue to more complex molecular transformations. By this way, RNA function can be fine-tuned at different levels depending on the type, location and target of the modification ^35^. Thus, certain RNA modifications are required for the proper processing, folding, and stability during RNA biogenesis, while others are crucial for RNA function in protein synthesis by modulating translation efficiency and fidelity ^36–39^. Noticeably, many of these RNA composition changes are not either static or stable ^40,41^. Instead, they are dynamic marks introduced and removed in a reversible process catalyzed by both epitranscriptomic “writer” and “eraser” enzymes, respectively, that can be potentially modulated in response to different cellular stress cues ^40,41^. However, this stress-induced reprogramming of RNA modifications and, specially, its likely connection with aneuploidy and derived therapeutic utility remain fully unexplored.

In an *in silico* bioinformatic work we have identified the *S*-adenosyl-methionine (SAM)-dependent RNA methyltransferase enzyme, TRMT61B, as a potential aneuploidy biomarker (to be published elsewhere). In the present work, we study in detail the connection of this molecule with highly aneuploid cancers and explore its potential therapeutic value. Despite it is encoded by the nuclear genome, TRMT61B is dominantly localized in the mitochondrial compartment where it catalyzes the methylation at N1 position of specific adenosine residues (referred as m^1^A) present in the three major mitochondrial RNA (mt-RNA) types ^42–44^. Although the molecular consequences of TRMT61B mediated methylation on RNA biology have received some attention, its biological function is still poorly understood ^42–44^. To this regard, very recent works suggest an involvement of this enzyme in protein synthesis regulation which is in agreement with the role of TRMT61B mtRNA targets as critical players of the mitochondrial translation machinery ^42–44^. However, a large body of research is required for addressing this issue in depth.

Here, our *in silico* and *in vitro* assays reveal a strong direct correlation between aneuploid degree and TRMT61B protein levels in both NCI60 cancer cell line collection and a more specific panel of human melanoma cell lines. Further, TRMT61B is found overexpressed in different tumor types compared to normal tissues and is positively associated with aneuploidy in human cancers. We also demonstrate that cancer cells with high levels of aneuploidy are greatly addicted to TRMT61B since its elimination by shRNA or CRISPR technology causes antiproliferative effects and negatively affects mitochondrial function. Finally, xenotransplant experiments performed in both immunodeficient mice and zebrafish embryos indicate a reduced tumorigenic capacity of TRMT61B deficient cells and reinforce the interest of this enzyme as a potential druggable target.

## Results

### TRMT61B as a novel cancer and aneuploidy biomarker

Considering the poor prognosis outcomes of cancer patients bearing highly aneuploid (ANE^high^) tumors and the clinical interest to explore specific properties that might allow ANE^high^ cancer cells to be efficiently killed, we aimed to discover novel molecular traits related to the ANE^high^ state. To this end, we took advantage of the publicly available data on the NCI-60 human cancer cell line panel, which has been molecularly, pharmacologically and karyotypically characterized with unparalleled depth and coverage, offering an interesting opportunity to carry out different genomic and gene expression association studies^45–52^. Using DNA ploidy information and four different structural and numerical karyotypic features, all the cell lines included in the panel were stratified following an unsupervised hierarchical clustering in four groups ranging from low to high aneuploidy levels (to be published elsewhere). To uncover those proteins strongly associated with aneuploidy, we compared the proteomic signatures ^49^ of the group with the highest levels of aneuploidy (ANE^high^) with the one with the lowest (ANE^low^). Taking as significant threshold FDR<0.15, the mitochondrial RNA methyltransferase TRMT61B was the unique protein found differentially altered, being overexpressed in the ANE^high^ group (**Fig. 1a**). Interestingly, the mRNA levels of TRMT61B were also significantly upregulated (adjusted P value of 0.011) among the ANE^high^ NCI60 cancer cell lines when compared with the ANE^low^ cell lines.

**Figure 1.**
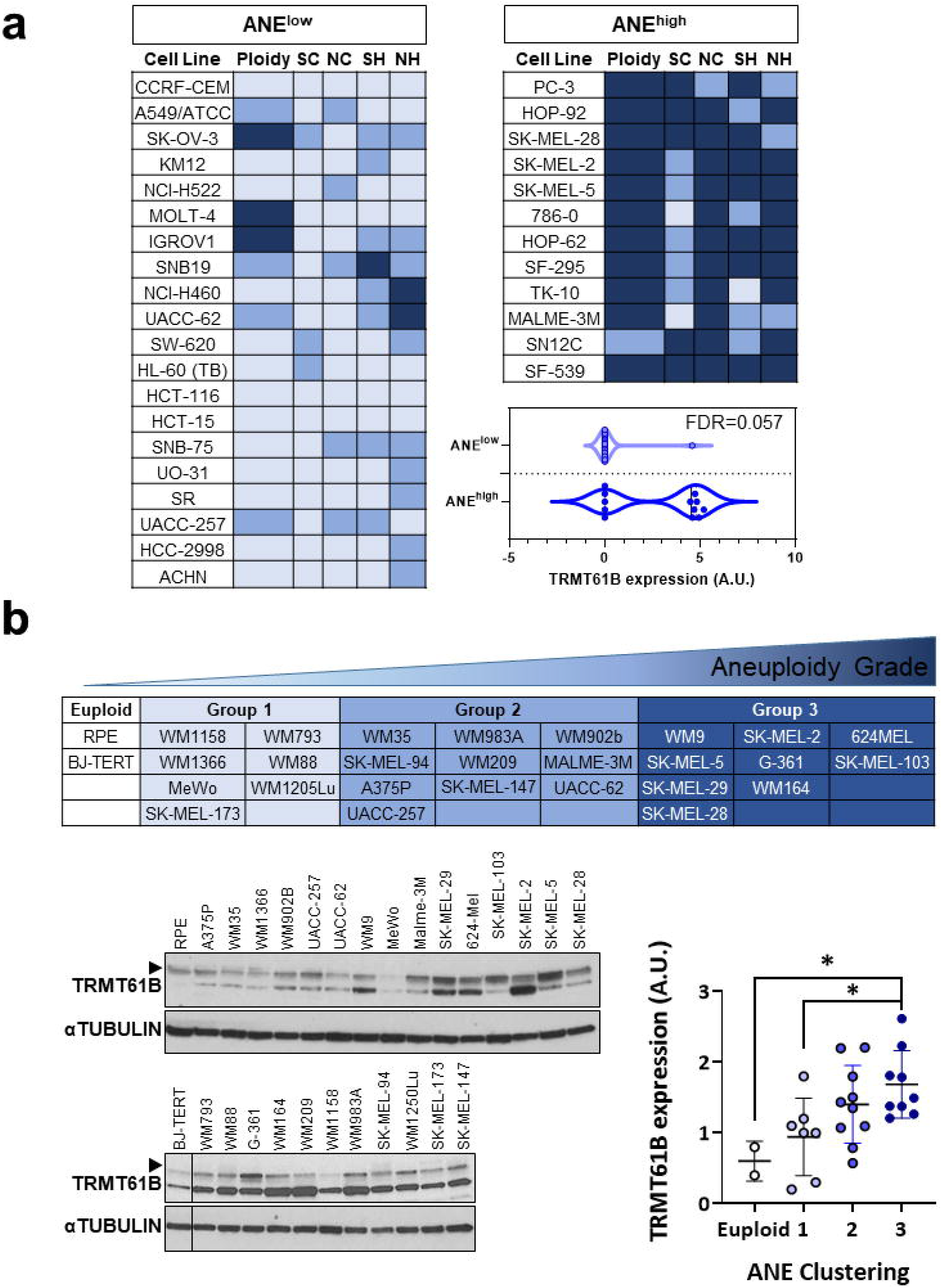
TRMT61B protein expression positively correlates with aneuploidy levels in human cell lines. **a** Following a bioinformatic analysis, tumor cell lines included in the NCI60 panel were divided according to different karyotypic features (SC, SH, NC, NH and ploidy) in four clusters ranging from low to high aneuploidy levels. Cell lines belonging to the most extreme groups in terms of aneuploid levels are listed. Proteome profile comparison between these two clusters allow us to identify *TRMT61B* gene, a mitochondrial RNA methyltransferase, whose protein expression levels were upregulated in ANE^high^ vs. ANE^low^ (FDR=0.057). SC: Structural Complexity; SH: Structural Heterogeneity; NC: Numerical Complexity; NH: Numerical Heterogeneity. **b** A collection of 26 human melanoma cell lines and 2 euploid controls were karyotypically characterized for some of the previous karyotypic parameters (SC, NC and ploidy), divided in four clusters (upper table) with null (euploid group) or increased rate of aneuploidy (1, 2 and 3) and finally analyzed by western blot for TRMT61B expression (bottom panels). As it is shown in the dotplot, group 3 displays a statistically significant upregulation of TRMT61B protein expression compared to euploid and 1 groups. Error bars represent standard deviation. \**P*< 0.05 (Student’s *t*-test; unpaired, 1-tailed).

To validate these *in silico* results, a large collection of 26 primary and metastatic melanoma cell lines, 6 of them shared with the NCI60 panel, was karyotypically characterized and examined by immunoblot for TRMT61B protein expression (**Fig. 1b**). Two non-tumoral cell lines were used as euploid controls and subjected to the same chromosomal and molecular characterization (**Fig. 1b**). Considering the structural and numerical chromosome alterations and their ploidy, cell lines were divided in four groups (euploid, 1, 2 and 3), being euploid and group 3 the ones with the lowest and highest aneuploidy levels, respectively (**Fig. 1b**). Quantification of TRMT61B expression showed a positive correlation between the aneuploidy grade and TRMT61B protein levels (**Fig. 1b**). Remarkably, only those comparisons between clusters exhibiting the most extreme values in terms of aneuploidy levels, 3 vs. euploid and 3 vs. 1, gave rise to statistically significant differences regarding TRMT61B expression with the highest average level corresponding to group 3 (**Fig. 1b**).

To explore in more detail the physiological relevance of all these findings, we carried out an extensive analysis of TRMT61B expression in different cancer types and contexts. First, we investigated TRMT61B mRNA expression *in silico* using publicly available information from different datasets. As shown in **Figure S1a** and **S1b**, TRMT61B mRNA expression is found frequently over-expressed in a wide range of cancer types and also in comparison with their corresponding healthy tissues. In a second analysis, we studied the potential connection between TRMT61B mRNA expression and aneuploidy scores using a TCGA tumor collection previously characterized by others in terms of aneuploidy levels (Table S1)^15^. Interestingly, a significant positive association between both variables was detected in 14 out of the 33 tumor types analyzed, whereas only for renal cell carcinomas (KIRC) higher TRMT61B expression was significantly associated with lower aneuploidy scores (**Fig. 2a**). To note, the significant associations accumulated in those cancers showing higher aneuploidy scores (**Fig. 2a**, towards the right part of the graph). Further, a positive and significant correlation between TRMT61B expression and aneuploidy grade could be detected when all the TCGA tumors were individually considered in the analysis (Fig. S1c). These bioinformatic results guided us to evaluate TRMT61B protein levels in two different tissue microarrays (TMAs) of cholangiocarcinoma (CCA) and melanoma (SKCM), that represent cancers with medium and high aneuploidy grades (**Fig. 2a**). The analysis of matched tumor-normal pairs revealed a clear upregulation of TRMT61B protein in the neoplastic lesions compared with healthy tissues in both cases (**Fig. 2b** and **2c**). However, no correlation of TRM61B expression with tumor stage was found for any of these tumor types (**Fig. 2b** and **2c**), which might indicate that TRMT61B over-expression is associated with tumor development but not with tumor progression.

**Figure 2.**
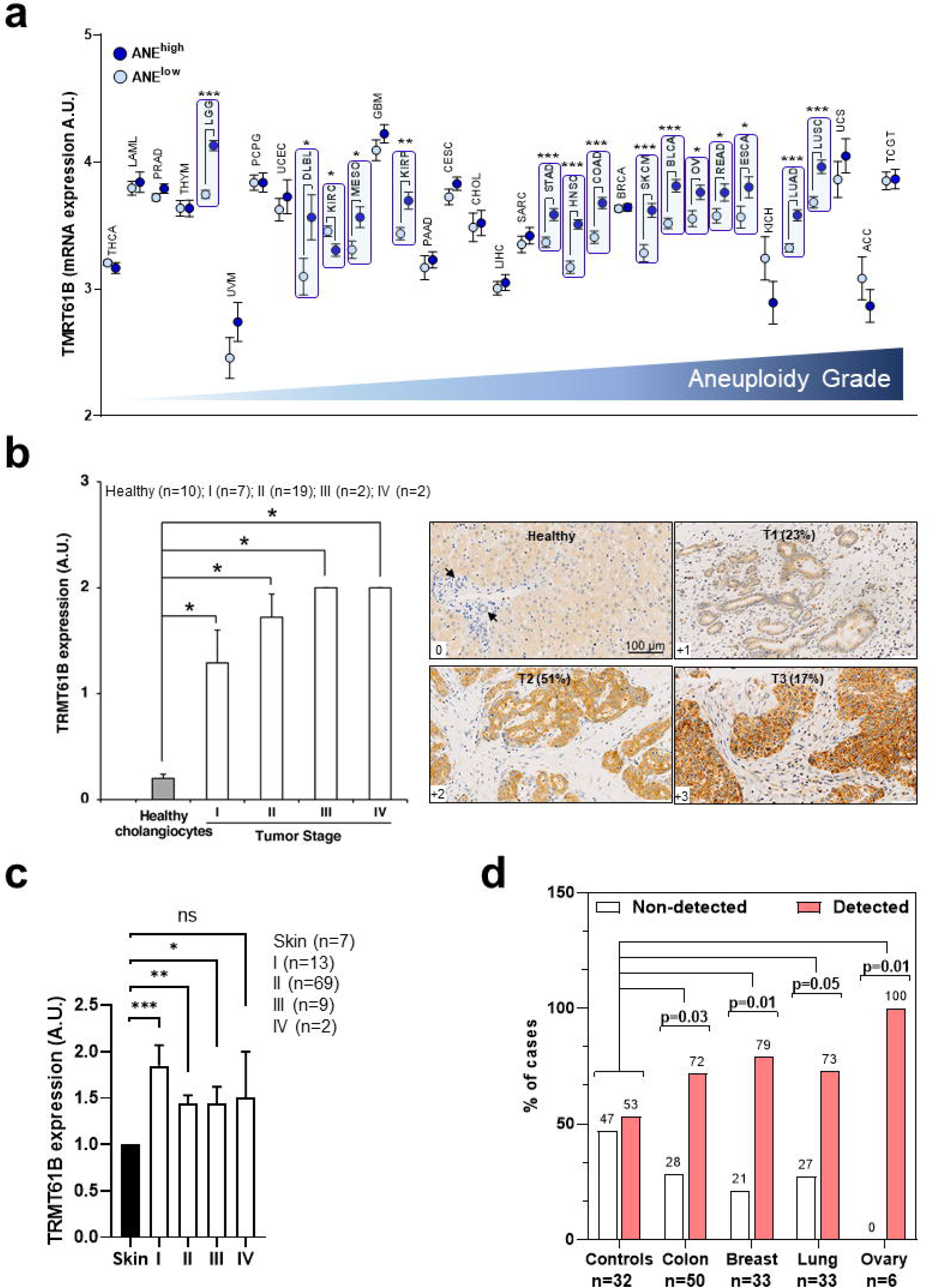
TRMT61B expression is positively associated with cancer and aneuploid levels in human cancers. **a** Examination of TRMT61B expression in several tumor types from the TCGA collection reveals increased mRNA levels of this RNA methyltransferase in cancer samples with higher aneuploidy scores. **b** TRMT61B levels are elevated in cholangiocarinoma (CCA) samples compared with healthy cholangiocytes without any positive association with tumor stage. Right panels depict representative immunohistochemistry images of cholangiocarcinomas (T1, T2 and T3) displaying different levels of TRMT61B expression (+1, +2, +3) and a healthy liver showing faint TRMT61B staining in the cytoplasm of hepatocytes and a very weak detection in cholangiocytes (near 0 in most of the cases) (arrows). In brackets, the percentage of samples exhibiting the indicated expression levels of TRMT61B is shown. Scale bars 100 μm. **c** Expression levels of TRMT61B protein is significantly elevated in melanoma human samples when compared with normal skin. No positive correlation is observed regarding tumor stage. **d** TRMT61B protein is detected in serum from cancer patients in a higher percentage of cases than in control serum. Error bars represent standard deviation. \**P*< 0.05 (Student’s *t*-test; unpaired, 1-tailed).

Finally, we explored the potential of TRMT61B as cancer biomarker in liquid biopsies by ELISA quantification of TRMT61B protein in serum samples of both healthy and tumor bearing patients. Although it was also found in 53% of the controls, TRMT61B detection accounted for more than 70% of the serums from colon, ovary, lung and breast cancer patients (**Fig. 2d**).

In summary, all these results indicate that TRMT61B expression is associated with higher aneuploidy and cancer development in a wide range of cell and tumor types.

### TRMT61B overexpression does not induce tumorigenesis nor alter chromosomal landscape

The fact that tumor cells showing increased aneuploidy tend to express augmented amounts of the RNA methyltransferase TRMT61B in sharp contrast to their normal and euploid counterparts, might be due to a positive cell selection during the tumorigenesis process based on aneuploidy grade and TRMT61B levels. However, it might also be possible that an initial abnormal up-regulation of TRMT61B induces aneuploidy and, subsequently, tumorigenesis. To test this last hypothesis, we decided to interrogate the potential oncogenic role of TRMT61B. Both euploid (RPE and BJ) and melanoma (MeWo) cell lines expressing reduced protein amounts of this RNA methyltransferase, were infected with a lentiviral construction encoding a flag-tagged, wild type TRMT61B form. Ectopic expression was confirmed by western blot and immunofluorescence analysis in the three cell-context examined (**Fig. S2**). As previously reported^42^, exogenous TRMT61B display a mitochondrial subcellular localization pattern, reinforcing the mitochondrial function originally attributed to this enzyme (**Fig. S2**). Using these cell models, we analyzed two main features associated with malignant transformation, cell proliferation and invasiveness. As shown in **Figure 3a**, elevated TRMT61B levels were associated with a slightly, though significant, decrease in cell proliferation rates of the three analyzed cell lines. On the other hand, invasiveness, another hallmark of transformation, was affected in the opposite way, mainly in the case of RPE cells (**Fig. 3b**). Importantly, when we examined the karyotype status of RPE and BJ cell lines overexpressing TRMT61B, we were unable to identify gross chromosomal rearrangements in comparison with their respective controls (**Fig. 3c**).

**Figure 3.**
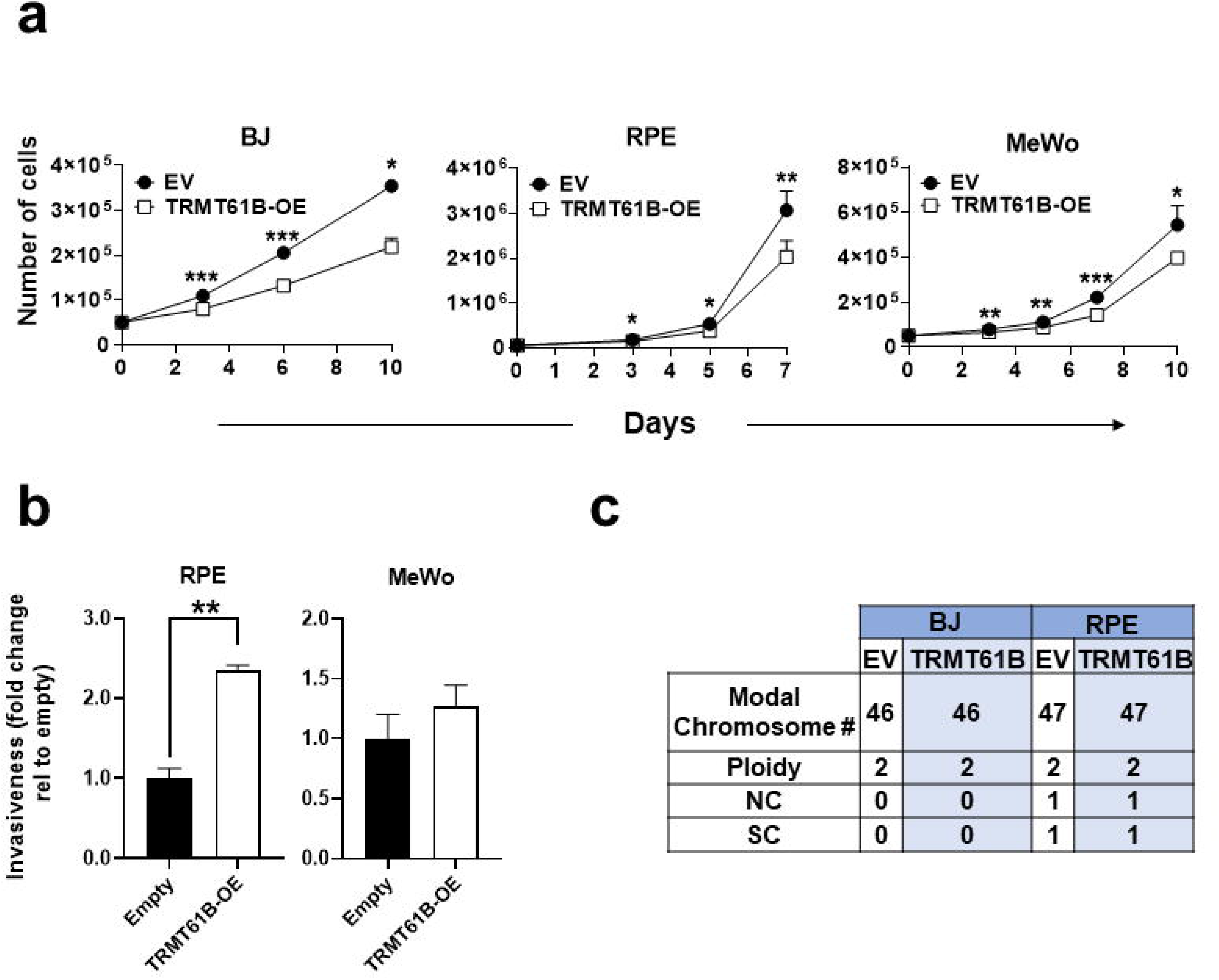
TRMT61B overexpression fails to produce karyotype abnormalities and causes milder effects on cell biology. **a** Proliferation of the indicated cell lines is slightly reduced after ectopic expression of TRMT61B (white squares) compared to controls (black circles). **b** *in vitro* evaluation by transwell assays of the invasive capacity of MeWo and RPE cells displaying basal (Empty) or increased levels (TRMT61B-OE) of TRMT61B. c Normal karyotype status of RPE and BJ cell lines regardless of TRMT61B expression levels. Error bars represent standard deviation. \**P*< 0.05; **0.001<*P*< 0.01; *** *P*< 0.001 (Student’s *t*-test; unpaired, 2-tailed).

According to these data, TRMT61B over-expression is not sufficient, per se, to induce, either aneuploidy or consistent tumor-associated features.

### Suppression of TRMT61B expression induces prominent antiproliferative effects in ANE^high^ cells

To determine the biological role of TRMT61B and the impact of its removal in a highly aneuploid context, we performed loss of function experiments by targeting TRMT61B expression via lentiviral shRNA interference (**Fig. S3**). Although our initial *in silico* results indicate that the connection between TRMT61B and aneuploidy is not restricted to any tumor type, we decided to focus on one unique cancer type, melanoma, which was well represented in the ANE^low^ and ANE^high^ groups of the NCI60 panel (**Fig. 1a**). To discard off-target effects, four different TRMT61B specific shRNAs and the corresponding scramble construction were analyzed in several melanoma cell lines. Since all of them efficiently diminish TRMT61B expression (**Fig. S4a**), they were indistinctly used along the different procedures. In a first set of experiments, three ANE^high^ (SK-MEL-2, SK-MEL-5 and SK-MEL-28) and two ANE^low^ (UACC-62 and UACC-257) melanoma cell lines included in the NCI60 collection together with two non-tumoral euploid cell lines (RPE and HBL-100), were efficiently silenced for TRMT61B expression and subsequently subjected to cell viability assays (**Fig. 4**). Our results showed that TRMT61B depletion reduces cell division capacity in all cases. However, ANE^high^ cells were more prominently affected since proliferation was completely halted even 16 days after lentiviral infection (**Fig. 4a**). Similar results were obtained with an independent collection of melanoma cell lines where those expressing increased amounts of TRMT61B were more sensitive to its elimination (**Fig. S4b**).

**Figure 4.**
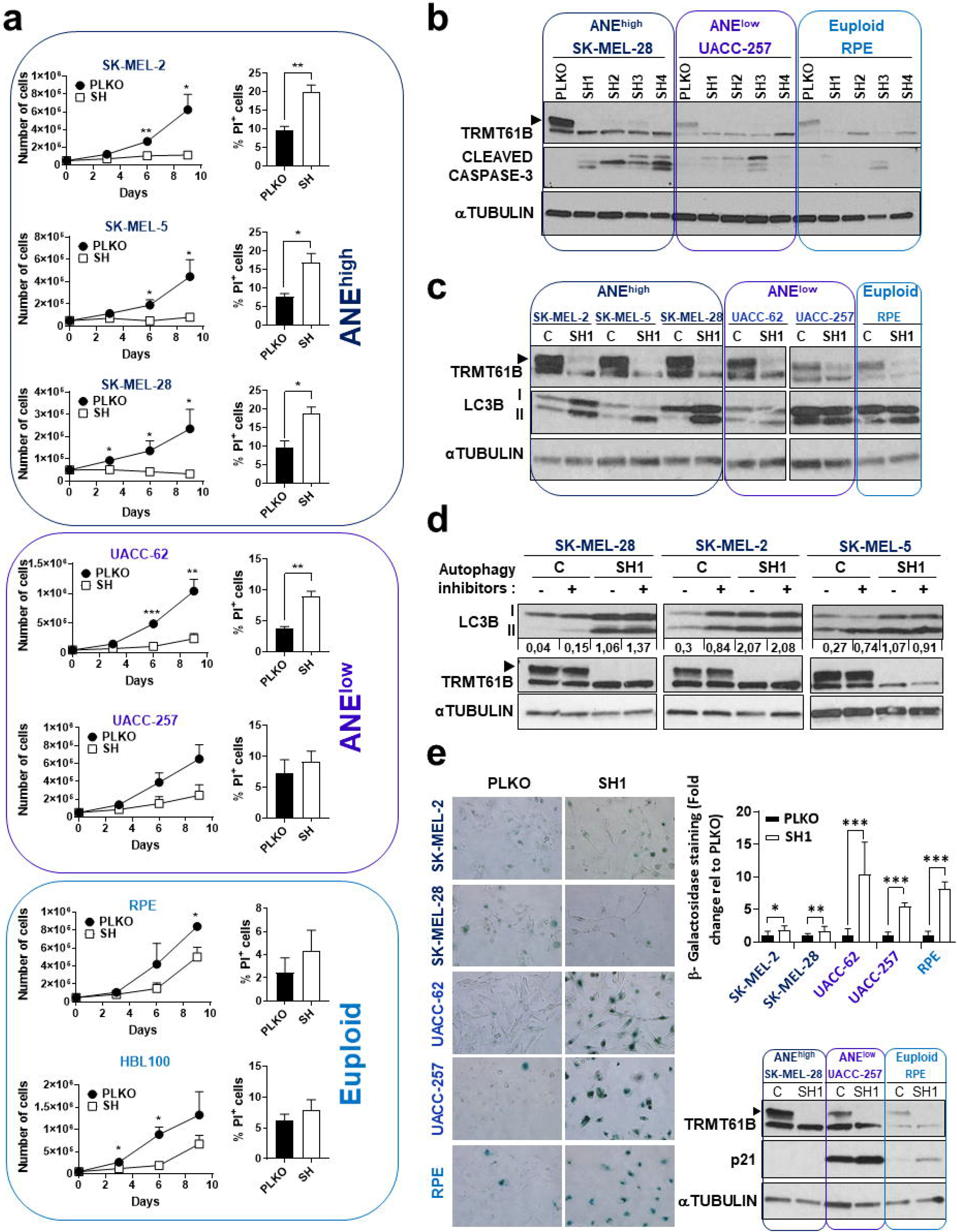
Antiproliferative effects induced by TRMT61B knock-down in cell lines with different aneuploid levels. **a** Proliferation ability (left graphs) of cells exhibiting different aneuploidy levels (euploid, low and high) according to the *in silico* analysis, is greatly compromised, especially in ANE^high^ cells, after TRMT61B interference with a concomitant apoptotic response (right columns). Results are the mean of 4 independent experiments performed with 3 different shRNAs (SH1, SH2 and SH4). Apoptosis is analyzed by PI staining and represents the average of 4 measurements taken at different time points (0, 3, 6 and 9 days after cell plating) during each of the experiments. **b** WB analysis of cleaved caspase-3 amounts in the indicated control and TRMT61B interfered cell lines reveals a strong induction in the ANE^high^ group. **c** WB analysis of the autophagy marker LC3BII in cell lines of the three ANE groups lacking or not TRMT61B expression. Similar results to the blot represented here were obtained from three independent experiments in which two different shRNAs (SH1 and SH4) were used for TRMT61B silencing. **d** The biological meaning of LC3BII induction detected in ANE^high^ cells after TRMT61B abrogation was interrogated by exposing both control and TRMT61B depleted cells to the autophagy inhibitory cocktail PepstatinA and E64d (5μg/ml, 7h). Measurement of the autophagy flux indicates a block of the autophagy cascade in the absence of TRMT61B. Three different experiments were performed with at least two of the cell lines. **e** β-galactosidase staining pictures (left panels) and quantification (right upper histogram) corresponding to control and melanoma cell lines showing normal or downregulated TRMT61B levels indicate a prominent senescence response in the ANE^low^ and euploid clusters. Correlation with p21 expression, a well-known senescence marker, is also shown (right lower panel). Two independent experiments were carried out in the case of SK-MEL-28, UACC-257 and RPE cell lines using two different shRNAs (SH1 and SH4). A representative experiment is depicted. Error bars represent standard error in **(a)** and standard deviation in **(e)**. \**P*< 0.05; **0.001<*P*< 0.01; *** *P*< 0.001 (Student’s *t*-test; unpaired, 2-tailed).

We then analyzed whether the absence of TRMT61B prevents cell growth by triggering apoptosis. To this end, propidium iodide (PI) staining of cultured cells and detection of cleaved caspase-3 in whole protein extracts were analyzed. Whereas euploid cells remain hardly affected, melanoma cells were characterized by a significant increase in the percentage of PI positive cells and the amounts of cleaved caspase-3 (**Fig. 4a, b** and **Fig. S4a, b**). As it occurred for the proliferation rates, the differences were more pronounced among ANE^high^ than ANE^low^ cancer cells. These results were also replicated in other set of melanoma cell lines (**Fig. S4a, b**).

Autophagy has been recently proposed as an important cell survival mechanism for CIN cells ^21^. Its protective role against apoptosis has been widely reported providing a basis for many pre-clinical and clinical studies where autophagy inhibitors are intended to increase tumor cell death when used in combination with other anti-cancer agents ^21,53–55^. Consequently, we sought to investigate the status of the autophagy process in response to TRMT61B depletion. Interestingly, the absence of this RNA methyltransferase leads to an accumulation of the autophagy marker LC3B-II in the three NCI-60 ANE^high^ cell lines (**Fig. 4c** and **Fig. S4c**, upper panel). On the contrary, LC3B-II levels remain unchanged in the ANE^low^ and euploid cell lines despite TRMT61B elimination (**Fig. 4c** and **Fig. S4c**, upper panel). In line of such evidence, analysis of other melanoma cell lines reveals a similar behavior after TRMT61B downregulation with increased amounts of LC3BII detected in most of the cell lines displaying higher aneuploidy and TRMT61B levels (**Fig. S4c**, lower panel). To properly understand the biological meaning of this finding, we treated both TRMT61B depleted and non-depleted ANE^high^ cells with the autophagy inhibitor cocktail E64d and pepstatin A known to disrupt the autophagy flux. As expected, blockage of the autophagy cascade yields a clear induction of LC3B-II in a TRMT61B proficient background (**Fig. 4d**, C lines). However, the treatment fails to produce a similar effect in a TRMT61B deficient context (**Fig. 4d**, SH1 lines), suggesting that TRMT61B depletion inhibits or slows down the autophagy process, at least in an ANE^high^ context.

Finally, considering that RPE, UACC257 and other ANE^low^ cell lines display unaltered LC3B-II amounts and PI indexes upon TRMT61B interference, we wonder whether other antiproliferative mechanisms are operating in parallel to produce the moderate anti-proliferative effect detected in the cell growth experiments (**Fig. 4a**). Therefore, we evaluated senescence response as a potential candidate able to be dysregulated by measuring β-galactosidase activity and p21 expression. Interestingly, both parameters were found significantly upregulated in ANE^low^ and immortal clusters once TRMT61B is silenced (**Fig. 4e**). Conversely, weaker, although still significant, differences were detected in the ANE^high^ cells, in the same direction.

Taking all these data together, we conclude that elimination of TRMT61B causes cytotoxic and cytostatic effects on aneuploidy cell survival by the occurrence of several antiproliferative mechanisms, whose intensity depends on the aneuploidy grade.

### Impaired mitochondrial morphology, respiration and translation due to TRMT61B deficiency

Given that TRMT61B exerts its function in the mitochondrial compartment ^42^, to understand the mechanisms underlying the effects induced by TRMT61B depletion, we studied at different levels the impact of TRMT61B downregulation in mitochondrial physiology. First, we analyzed the architecture of this organelle using electron microscopy. As shown in **Figure S5a**, mitochondria appear partially devoid of crests in cells infected with shRNAs specific for TRMT61B. Since ATP generation through oxidative phosphorylation (OXPHOS) powered by the electron transport chain complex (ETC) is one of the most prominent mitochondrial functions, we aimed to carefully analyzed the consequences of TRMT61B loss in the respiration metabolism ^56^. For this purpose, we measured the respiratory activity of ANE^high^, ANE^low^ and euploid cells lacking or not TRMT61B by using a Seahorse XF96 analyzer. As shown in **Figure 5a**, mitochondrial respiration of two out of three 3 ANE^high^ and one out of two ANE^low^ cell lines was greatly compromised upon TRMT61B depletion. A similar trend but not statistically significant could be observed in the case of the SK-MEL-2 cell line belonging to the ANE^high^ group (**Fig. 5a**). No differences were detected for UAC-257 (ANE^low^) and RPE (euploid) cells.

**Figure 5.**
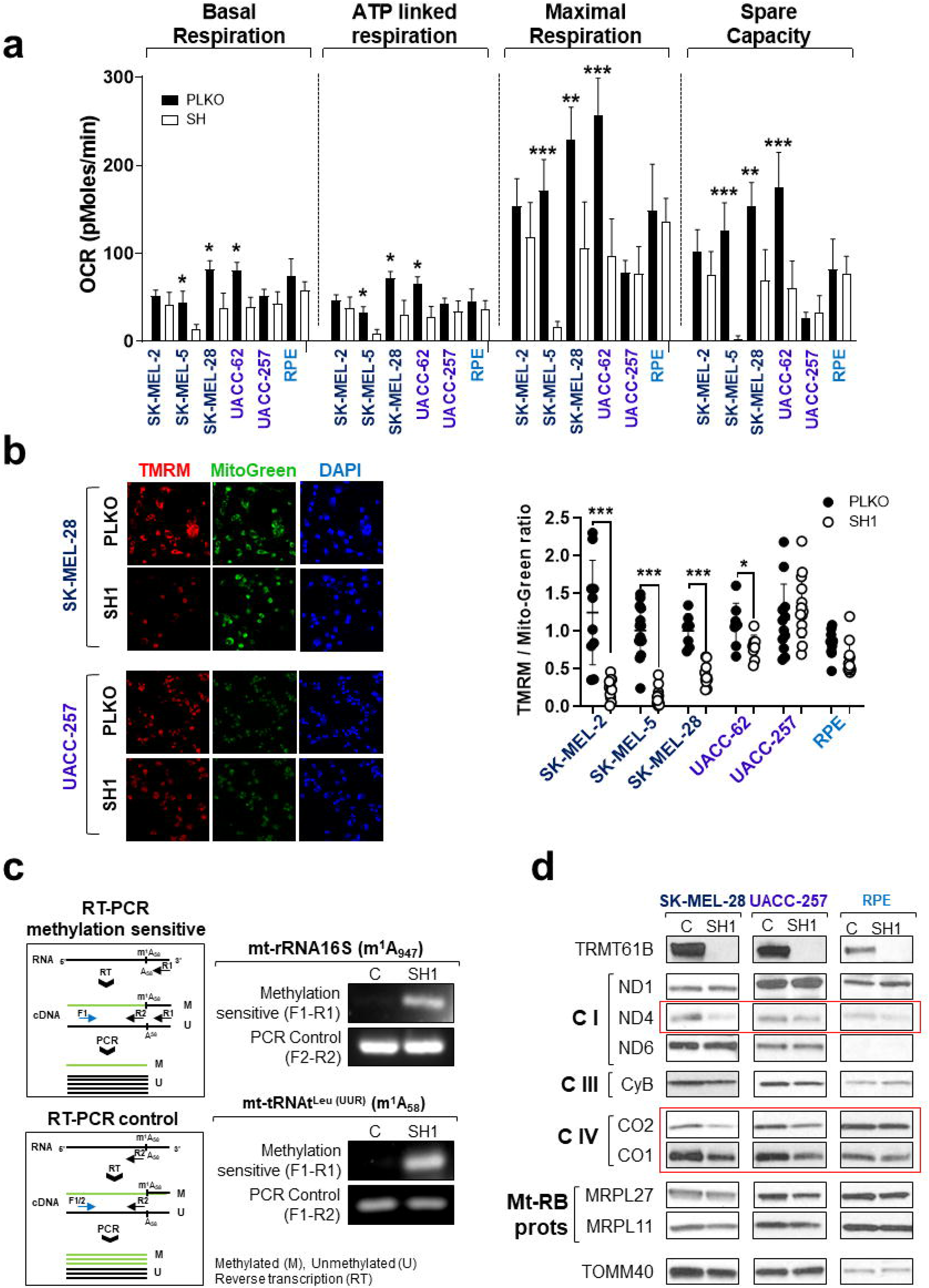
TRMT61B is required for mitochondrial functionality. **a** Seahorse analysis of OXPHOS (oxidative phosphorylation) reveals a statistically significant reduction of ATP-linked, basal and maximal respiration as well as spare capacity (difference between maximal and basal respiration) in two out of three ANE^high^ (SK-MEL-28 and SK-MEL-5) and one out of two ANE^low^(UACC-62) cell lines, once TRMT61B is eliminated, suggesting that this RNA methyltransferase is required for a proper stress response. Continuous OCR values (pmoles/min) are shown. Mitochondrial functions were analyzed as explained in Materials and Methods. Five independent experiments were carried out between five and seven days after TRMT61B silencing using two different shRNAs (SH1 and SH4). Error bars represent standard error. \**P*< 0.05; **0.001<*P*< 0.01; *** *P*< 0.001 (Student’s *t*-test; unpaired, 2-tailed). **b** In the same group of cells also including SK-MEL-2 cell line, TMRM staining, a fluorescent probe accumulated in active mitochondria, reveals a clear loss of mitochondrial membrane potential upon TRMT61B downregulation, since a diminished signal is detected after quantification (right dotplot diagram). Conversely, RPE and UACC-257 cells do not display significant differences in TMRM staining independently of TRMT61B status. Representative fluorescence images are depicted showing TMRM (red), Mitotracker (green) and DAPI (blue). Error bars represent standard deviation. \**P*< 0.05; *** *P*< 0.001 (Student’s *t*-test; unpaired, 2-tailed). **c** PCR strategy to determine methylation levels (left upper scheme) of the indicated mt-RNA types based on the ability of N1-methyladenosine (m^1^A) to block reverse transcription. Amplification levels of the resultant cDNA by semi-quantitative PCR allow to indirectly estimate the presence of the desired methylation mark. As expected, TRMT61B silencing reduces specific m^1^A formation (residues in brackets) in two known mt-RNA targets according to the detected PCR product. A second PCR strategy (left lower scheme) insensitive to m^1^A methylation is also described and used as a loading control. **d** Protein expression levels of several mitochondrial encoded genes that are part of the electron transport chain (Complexes I, III and IV) as well as nuclear encoded mitoribosomal proteins (Mt-Rb), were analyzed by western blot in mitochondrial protein extracts derived from the indicated cell lines expressing basal or reduced levels of TRMT61B. The mitochondrial marker TOMM40 was used as a loading control since its expression is not expected to be altered. Genes highlighted in red are the most affected when TRMT61B is removed.

This substantial reduction in the oxygen consumption rate (OCR) after TRMT61B downregulation is reflected in a decreased basal respiration, ATP-linked respiration and especially, in both maximal and spare capacity respiration, strong indicators of mitochondrial dysfunction (**Fig. 5a**). Importantly, no glycolytic compensation could be detected in any of the cases (**Fig. S5b**). We next measured the mitochondrial membrane potential (ΔΨ_m_) as it represents a good indicator of the mitochondrial health status. Cells were loaded with tetramethylrhodamine methyl ester (TMRM) and live imaging was used to estimate ΔΨ_m_. In agreement with previous bioenergetic results, we found that TRMT61B deficient ANE^high^ cells and UACC-62 cell line (ANE^low^) show a dramatic decrease in the ΔΨ_m_ compared to their respective controls according to the significant decay in the TMRM/Mitotracker-greeen ratio (**Fig. 5b**). In parallel to seahorse stress test, no signs of mitochondrial depolarization could be detected in both UACC-257 and RPE cell lines devoid of TRMT61B (**Fig.4b**).

We next asked whether lack of TRMT61B affects mitochondrial gene expression since the 13 protein-coding genes encoded by the mitochondrial genome are involved in the formation of the ECT complex, at least in part, with the aid of several TRMT61B mt-RNA targets^42–44,57^. Thus, 7-10 days after TRMT61B silencing mitochondrial fractionation was performed for one representative cell line of each of the ANE groups and mitochondrial RNA and proteins were studied. As expected, m^1^A marks in two of the best characterized TRMT61B RNA substrates, mt-RNA 16S and mt-RNA^leu 42,43^, were greatly diminished in a TRMT61B depleted context (**Fig. 5c**). On the other hand, western blot analysis revealed significant reduction in the expression levels of some mitochondrial encoded genes including CO2, CO1 and ND4. This reduction tends to be slightly more prominent in the case of the ANE^high^ and ANE^low^ cell lines defective for TRMT61B expression (**Fig. 5d**).

Altogether these data show that TRMT61B is necessary for efficient mitochondrial genome expression linking TRMT61B downregulation to OXPHOS deficiency. This process is especially relevant in ANE^high^ cells, which could require a larger respiration capacity.

### TRMT61B inhibition decreases *in vitro* cell invasion and *in vivo* tumorigenesis

Our results suggest that by negatively affecting mitochondrial function and cell viability, TRMT61B abrogation could be an antitumoral target with significant value in a highly aneuploidy context. To address this hypothesis, we sought to gain further insights on the effect of TRMT61B downregulation in tumorigenesis both *in vitro* and *in vivo*. First, we analyzed the invasive capacity of control and TRMT61B deficient cells from the three aneuploidy groups. Regardless of the aneuploidy grade, downregulation of TRMT61B significantly diminishes cell mobility through a matrigel basement membrane matrix suggesting a potential negative impact on metastatic capability (**Fig. 6a**). Going one step further, we examined the *in vivo* effect of TRMT61B depletion on tumorigenesis by subcutaneous injection into immunocompromised mice of SK-MEL-103 melanoma cells transduced with a doxycycline inducible TRMT61B specific shRNA construction (**Fig. S6a**). This inducible model of TRMT61B knockdown was previously characterized *in vitro* by western blot and cell proliferation assay showing a significant antiproliferative phenotype upon doxycycline mediated TRMT61B silencing (**Fig. S6b**). Remarkably, *in vivo* administration of doxycycline after tumor onset significantly attenuates tumor growth despite the incomplete interference of TRMT61B expression (Fig. 6b). Further, tumors developed by doxycycline treated animals displayed decreased Ki67 and increased cleaved caspase-3 staining, which strengthens the antitumoral effect mediated by TRMT61B silencing (Fig.6b, right panels).

**Figure 6.**
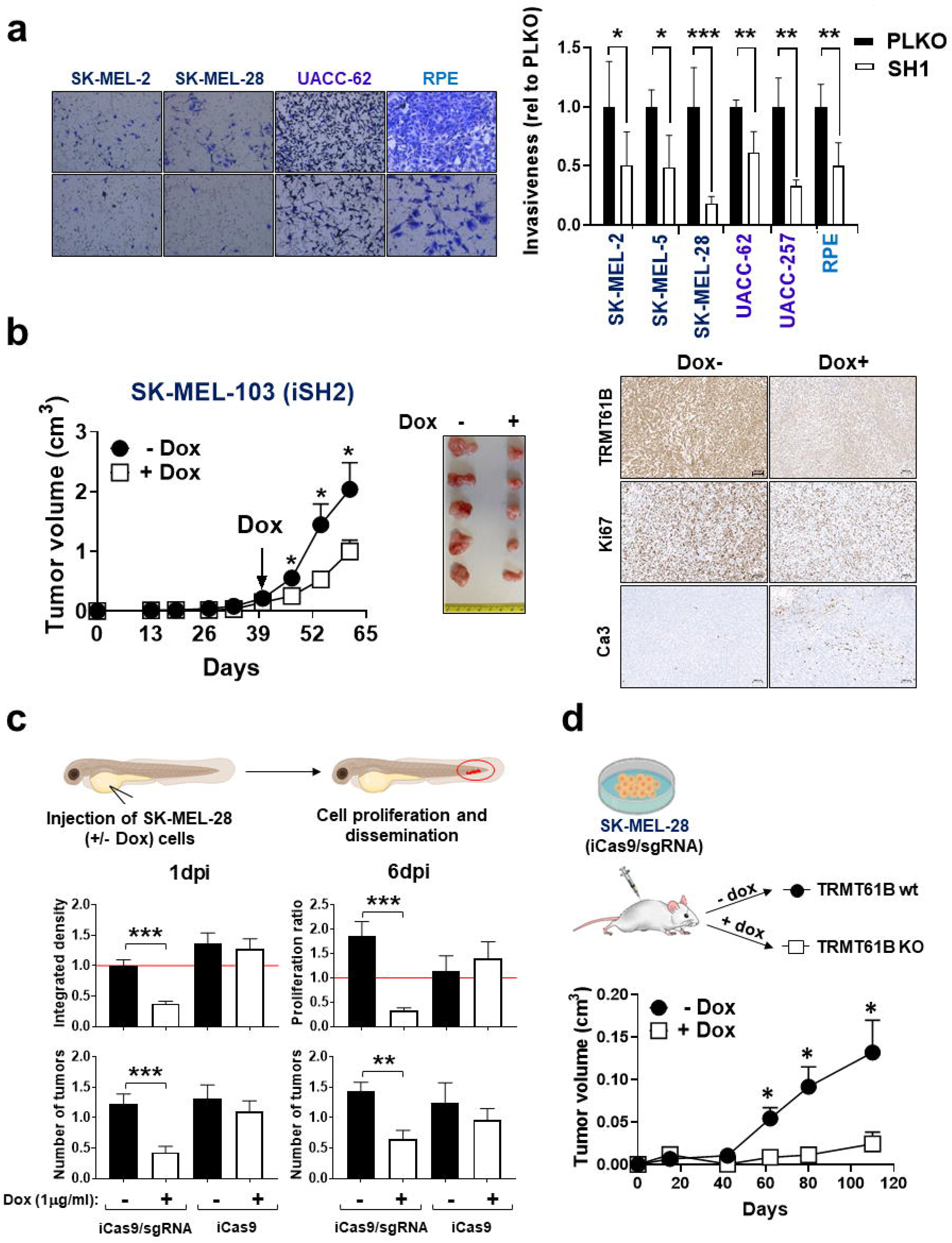
TRMT61B abrogation diminishes tumorigenic potential. **a** ANE^high^, ANE^low^ and euploid cells lentivirally infected with a TRMT61B specific shRNA (SH1) exhibit impaired invasive capacity through matrigel coated inserts compared with scramble shRNA transduced counterparts. Data are normalized against control values and two different experiments were carried out in triplicate. One representative experiment is shown. Error bars represent standard deviation. \**P*< 0.05; **0.001<*P*< 0.01; *** *P*< 0.001 (Student’s *t*-test; unpaired, 2-tailed). **b** Tumor growth in Nod/scid mice injected with SK-MEL-103 cells transduced with a doxycycline inducible shRNA targeting TRMT61B expression. Monitoring tumor volume by external caliper indicates a statistically significant reduction in tumor growth after doxycycline administration (left graph). Two different experiments were carried out with 10 mice per group (doxycycline treated and untreated). Error bars represent standard error. \**P*< 0.05 (Student’s *t*-test; unpaired, 1-tailed). Representative macroscopic images and immunohistochemical staining of TRMT61B, Ki67 and cleaved caspase-3 in tumor sections from each mouse cohort are represented (right). Scale bars 100μm. **c** Integrated density (1dpi) and proliferation ratio (6dpi) of SK-MEL-28 cells (both Cas9/sgRNA and Cas9 alone) injected into the circulation of the zebrafish embryos. The red line at 6dpi represents a threshold below which cell proliferation stops. Integrated density at 1dpi is a total measure of the fluorescence of the fish tails. Average of tumor number present in the tail of each condition tested at 1 and 6dpi is also indicated (lower left histograms). n_replica_=20 embryos/condition (3 replica), n_total_=60 embryos/condition; Error bars represent standard error. \**P*< 0.05 (Student’s *t*-test; unpaired, 1-tailed). **d** SK-MEL-28 cells containing both CRISPR/Cas9 elements were subcutaneously injected in nude mice being treated or not with doxycycline from the very beginning to schedule a preventive strategy. As it is shown in the graph, doxycycline administration causes a statistically significant tumor growth retardation. One experiment was performed with 12 animals per mouse cohort. Error bars represent standard error. \**P*< 0.05 (Student’s *t*-test; unpaired, 1-tailed).

As a complementary approach, we performed a doxycycline inducible CRISPR/Cas9-mediated gene editing approach to abolish TRMT61B expression in SK-MEL-28 cells (**Fig. S6c**). Western blot analysis revealed the presence of minimal TRMT61B amounts in doxycycline treated cells harboring both CRISPR/Cas9 elements (iCas9/sgRNA+Dox) compared to non-treated counterparts (iCas9/sgRNA-Dox) and Cas9 controls subjected (iCas9+Dox) or not (iCas9-Dox) to doxycycline treatment (**Fig. S6c**). Remarkably, iCas9/sgRNA+Dox cells displayed reduced proliferation rates, LC3B-II accumulation and slight cleaved caspase-3 induction (**Fig. S6c**). As expected, iCas9/sgRNA cells grown in a doxycycline-free medium as well as iCas9 ^+^Dox and iCas9-Dox cells, did not show either proliferative defects or LC3B-II changes (**Fig. S6c**). Further, specific methylation marks and expression of several mitochondrial protein coding genes were diminished in iCas9/sgRNA+Dox cells compared to controls (**Fig. S7**). Next, both iCas9/sgRNA and iCas9 cells were injected in zebrafish embryos to monitor their tumorigenic behavior after being previously subjected or not to an *in vitro* 10-day doxycycline treatment (**Fig. 6c**). Zebrafish embryos harboring pretreated iCas9/sgRNA+Dox cells, display reduced tumor burden and fluorescence signal at the end of the experiment (6dpi) in the continuous presence of doxycycline, while embryos bearing control cells (iCas9/sgRNA-Dox, iCas9-Dox and iCas9+Dox) maintain or increase the initial values for both parameters (**Fig. 6c**). In parallel, we also carry out similar *in vivo* experiments subcutaneously injecting iCas9/sgRNA SK-MEL-28 cells in immunodeficient mice (**Fig. 6d**). Once inoculated, two different animal cohorts were established receiving or not doxycycline in the drinking water from the initial stages (**Fig. 6d**). Similar to the zebrafish experiments, doxycycline administration causes a conspicuous tumor growth retardation (**Fig. 6d**).

These results indicate that TRMT61B function is critical to melanoma progression and highlights its utility as a druggable target for upcoming anticancer interventions.

## Discussion

Aneuploid is a characteristic feature of the great majority of cancer cells distinguishing them from their normal counterparts ^5,58^. Chromosome copy number and structural changes affect large cohorts of genes (including oncogenes, tumor suppressor genes, DNA repair genes, metabolic genes) that characterize the driving force behind cancer development ^59–61^. Moreover, aneuploidy and CIN are considered one of the main genetic diversification process causing intratumoral heterogeneity, metastasis and resistance to chemotherapeutics and, therefore, poor patient prognosis, in the context of cancer disease ^5,62^. However, before reaching this point, aneuploid cells need to overcome different stresses imposed by gene dosage imbalance and protein stoichiometry changes ^63^. It is widely accepted that aberrant genes connected to CIN and aneuploidy normally function in pathways involved in DNA replication and repair as well as in chromosome dynamics ^58,64^. Nevertheless, cross-species studies have allowed to successfully identify additional gene functions within less intuitive pathways that are not directly related to the processes listed above such as, tRNA synthesis, lipid synthesis or proteasome function ^65^. It is tempting to speculate that probably some of these new molecular players and others not yet identified might critically participate in the cellular, stress responses required for CIN/aneuploidy tolerance. Thus, the discovery and functional characterization of new genes associated with CIN/aneuploidy will help to find new anti-cancer treatment strategies. Our research efforts focused on identifying novel aneuploidy associated genes allowed us to uncover TRMT61B. The protein expression of this almost unknown RNA modifier correlates with the extent of chromosomal alterations in a large collection of human cancer cell lines included in the NCI60 panel. This interesting finding was experimentally and computationally validated in several human melanoma cell lines and TCGA tumor types. The TCGA validation strategy is especially relevant since despite having been considered TRMT61B expression in terms of mRNA levels and used different criteria for aneuploidy measurement, a positive correlation between both parameters could be found, which reinforces the potential of TRMT61B as an aneuploidy biomarker in a wide spectrum of tumor types. Further, immunohistochemistry analysis of TRMT61B in various tumor types and the corresponding unaffected samples, allowed us to correlate increased TRMT61B protein expression with cancer pathogenesis (Fig. **2**). Importantly, this correlation is not universal, being only found in a set of tumor types for which, so far, we have not found a common feature. Further detailed studies are needed to precisely determine for what tumors and stages TRMT61B expression can be used as a cancer and/or aneuploidy marker.

On the other hand, supraphysiological expression of TRMT61B failed to induce detectable karyotype changes indicating that TRMT61B is not a driving force behind aneuploidy but rather an adaptive or tolerance inducing molecule to an, *a priori*, unfavorable situation. In fact, this is in good correlation with many studies that report a role of RNA modifications in different cancer hallmarks like survival, proliferation, self-renewal, differentiation, stress adaptation and invasion instead of working as cancer drivers ^66^. Noticeably, as far as we know, it is the first time a study of such magnitude is conducted for TRMT61B given that previous works only established indirect or poorly detailed correlations in a very limited pathological contexts ^67,68^. This contrasts with the increasing data supporting the oncogenic and tumor suppressor roles of other RNA epigenetic players required for the regulation of better characterized, and in some cases more abundant, RNA modifications like m^6^A (N^6^-methyladenosine) or m^5^C (5-methylcytosine) ^66,69^.

Collectively, our data might indicate that TRMT61B upregulation reflects a gene addiction scenario able to be therapeutically exploited, as it has been shown in several studies where these RNA-modifying enzymes contribute to maintain cell proliferation and tumor progression ^69^. In line with this hypothesis, TRMT61B loss-of-function experiments performed using shRNA and CRISPR/Cas9 technologies induce an antiproliferative phenotype due to the co-occurrence of several cytotoxic and cytostatic responses acting depending on aneuploidy levels. The explanation for this limited proliferation relies on TRMT61B biological function that has remained under-investigated to date. This enzyme is one of the few RNA methyltransferases able to catalyze methylation of tRNAs, mRNAs and rRNAs, key elements of the translation machinery ^42–44^. Spite of being encoded by the nuclear genome, TRMT61B exerts its function in the mitochondrial compartment where works as a writer enzyme installing m^1^A modifications in specific positions of its RNA targets. Importantly, TRMT61B belongs to a reversible and dynamic regulation system for the proper m^1^A burden composed by several nuclear and mitochondrial writer (TRMT10C, TRMT6/61A), eraser (ALKBH1 and ALKBH3) and reader (YTHDF1-3, YTHDC1) proteins found to be dysregulated in several tumor types but with limited information at the moment ^70–72^. It has been shown that the positive charge carried out by m^1^A plays a critical role in maintaining the correct tRNA tertiary structure and function, potentially affecting the binding of the elongation factor that delivers tRNA to ribosome, which represents an emerging layer in the regulation of gene expression ^73–75^. In fact, TRMT61B depleted cells exhibit reduced expression levels of several mitochondrial encoded genes, components of the ETC, in parallel with a severe reduction of the m^1^A epitranscriptomic marks in at least two of the best characterized TRMT61B RNA targets. Surprisingly, although the magnitude of the expression change detected in the affected mitochondrial genes is quite similar between ANE^high^ and ANE^low^ cells, the latter and the non-tumoral cells display, in most of the cases, unaltered cell respiration and mitochondrial membrane potential upon TRMT61B silencing. This lack of effect in normal cells is well exemplified by different studies reporting the specific susceptibility of cancer cells to the genetic inhibition of enzymes modifying wobble uridine 34 (U34) tRNAs which seem to be critical for the efficient translation of several mRNA candidates encoding pro-invasive proteins ^76^. In the cases reported here, it could be possible that energy requirements and TRMT61B dependence are higher for those cells with an elevated aneuploidy degree since this status demands a more efficient activity (and expression of its components) of the ETC to generate ATP in an oxygen dependent manner. In contrast to the historical assumption about aerobic glycolysis as the major cancer energy source, many tumors reprogram their metabolism increasing mitochondrial activity to properly respond to the newly energetic and anabolic needs imposed by several stressors like CIN/aneuploidy ^28,29,77^. In the particular case of melanoma, although the consensus in the field also attributes to glycolysis a predominant metabolic role, mainly in a BRAFV600 context, emerging data shows that mitochondrial energy metabolism also coexists and could represent a promising therapeutic target^78–80^. Interestingly, similar to respiratory ATP depletion mediated by TRMT61B loss, it has been shown that chemical inhibition of OXPHOS reduces tumor growth and metastasis^81,82^.

Besides ATP production, other key functions of the cells, such as redox balance maintenance, programmed cell death or even autophagy, among others, require the critical participation of mitochondria ^83,84^. In this sense, it has been recently shown that CIN cells display a robust activation of autophagy to manage metabolic and oxidative stresses within tolerable levels ^21^. Further, inhibition of autophagy leads to massive apoptosis, DNA damage and increased ROS production ^21^. Additionally, complete deletion of Atg7 or Atg5, essential autophagy genes, negatively affects tumor growth in Ras- or Raf- driven tumors, suggesting a role of autophagy as a tumor promoting cellular pathway ^85–88^. In our experiments, TRMT61B depletion in ANE^high^ context hampers or delay autophagy flux and causes apoptosis. However, whether TRMT61B is directly or indirectly involved in autophagy regulation remains unclear. Shedding light on this aspect, a recent study performed in OXPHOS deficient fibroblasts harboring well characterized mutations in nuclear and mitochondrial OXPHOS genes, show a partial blockage of the autophagy cascade suggesting a connection between both processes ^89^. Further, it has been reported by Thomas and co-workers that phenformin treatment, an inhibitor of mitochondrial complex I, or genetic defects in complex I, suppress autophagy induced by mTOR inhibitors ^90^. Interestingly, several m^1^A regulator genes have been associated with the mTOR pathway in different gastrointestinal tumors ^70^. Importantly, inhibitors of other mitochondrial respiration complexes appear to be also potent autophagy disruptors ^91^. Considering that TRMT61B downregulation diminishes the expression of various protein components of at least complex I and IV, an indirect link between TRMT61B and autophagy can be proposed. Nevertheless, we cannot exclude the possibility that autophagy was a direct TRMT61B target whose inhibition could have a negative impact on mitochondrial metabolism, as it has been recently reported in acute myeloid leukemia^92^. Consequently, a complex network of interactions might be simultaneously working in a TRMT61B dependent manner and being influenced each other. Ideally, these pending questions and all the indicated findings should be addressed in a more physiological context, being mouse models the preferred experimental system. However, the absence of TRMT61B expression in murine cells as well as some of their molecular targets ^43^ has prevented us from approaching this issue with such strategy. Humanized mouse models or selection of alternative organisms might be valuable tools to get this ambitious objective.

Here, we have demonstrated that, in a highly aneuploidy cancer context, TRMT61B critically regulates mitochondrial function and its defective expression leads, in a not yet determined order, to autophagy impairment and translation defects of proteins involved in ETC and ATP synthesis. The resulting energy decay and accumulation of dysfunctional intracellular components like aberrant mitochondria, might inevitably push aneuploid cancer cells to their tolerance threshold threatening their survival. Therefore, targeting this epitranscriptomic modulator could be an effective way to perturb cellular stress responses critically required for aneuploid cell homeostasis and open novel avenues to design antitumoral treatments especially driven against more aggressive neoplasias.

## Methods

### Ethics statement

All mouse studies were performed in compliance with the institutional guidelines for the welfare of experimental animals approved by the Use Committee for Animal Care from Instituto de Salud Carlos III (ISCIII) and La Comunidad de Madrid and in accordance with the guidelines for Ethical Conduct in the Care and Use of Animals as stated in The International Guiding Principles for Biomedical Research involving Animals, developed by the Council for International Organizations of Medical Sciences (CIOMS).

All the procedures used in the zebrafish experiments as well as fish care were performed in agreement with the Animal Care and Use Committee of the University of Santiago de Compostela and the standard protocols of Spain (Directive 2012-63-UE). At the final point of the experiments, zebrafish embryos were euthanized by tricaine overdose.

Liver samples were obtained from patients having undergone surgical resection at the University Hospital of Salamanca, Spain. The study was approved by the Ethics Committee for Clinical Research of Salamanca (July 18, 2018) and informed written consent for the samples to be used for biomedical research was obtained from each patient.

Plasma samples were obtained from the biobank of the Hospital Clínico San Carlos or provided by Hospital Universitari de Sant Joan (Tarragona, Spain) after approval of the Ethical Review Boards of these institutions. All subjects in the study gave their written informed consent to participate and all experiments were performed in accordance with relevant guidelines and regulations.

### Cell lines and culture conditions

MeWo (A), A375P (A), 624-MEL (A) and WM35 (A) were a gift from Amparo Cano’s Laboratory. SK-MEL-2 (A), SK-MEL-5 (A), SK-MEL-28 (A), SK-MEL-173, SK-MEL-147, SK-MEL-103, SK-MEL-29, UACC-62(A), UACC-257 (A), WM1158, WM1205Lu, WM793, WM1366 (A), WM88, WM902B (A), WM209, G-361, Malme-3M(A), WM983A, WM9 (A) and WM164 (A) were kindly provided by Marisol Soengas’ group. Finally, BJ-TERT and RPE-TERT (A) were obtained from Marcos Malumbres’ laboratory. Since most of them were already authenticated by previous users, we confirmed the identity of those cell lines (with an “A” in brackets) more relevant for the study by STR DNA profiling. All cell lines were routinely tested for mycoplasma contamination, grown in DMEM medium supplemented with 10% FBS (Fetal Bovine Serum) and 1% penicillin-streptomycin and cultured at 37°C under a 5% CO_2_ atmosphere. Cells subjected to doxycycline treatment were exposed to a final concentration of 1-2 μg/ml during a minimum period of 4 days.

### Karyotype analysis

Exponentially growing cells were treated with 0.5mg/ml of the microtubule-depolymerizing drug colchicine for 4-6h, then centrifuged with a table-top centrifuge, swollen in 40% medium/60% tap water for 15 min at 37 ºC, fixed with Carnoy solution (75% methanol and 25% acetic acid) and spread on a glass slide. Metaphase chromosomes were GTG-Banded by a conventional trypsin-Giemsa technique and karyotypes (20-40 metaphases per cell line) were analyzed using a capture and analysis Cytogenetic software (Citovysion Version 7.7, Leica Biosystems Richmont, inc.).

### Aneuploidy based clustering of melanoma cell lines

Numerical (NC) and structural chromosome complexity (SC) values were obtained for each cell line, ranked and divided in three groups with high, intermediate and low NC and SC levels that were transformed in 1, 2 and 3 numbers, respectively. These processed values together with the ploidy content (ranging from 2 to 5) were aggregated and aneuploidy scores were assigned to each cell line. Finally, cell lines were classified in three clusters where groups 1 and 3 display the lowest and highest aneuploidy scores, respectively. For the analysis of the aforementioned karyotypic features, we follow the criteria described by Roschke and coworkers based on the ISCN convention^50^.

### Pan-Cancer Computational analyses

In the TCGA database, TRMT61B expression levels were calculated as log_2_(TPM + 1) transformed counts for 9131 tumor samples (belonging to 33 different cancer types) profiled by RNAseq. For these same tumor samples, aneuploidy scores were called from Taylor AM et al., 2018 ^15^. Tumors were subdivided into top and bottom quartiles based on their aneuploidy scores and compared for TRMT61B expression. Processed TCGA expression data were kindly provided by CNIO Bioinformatic Unit (see Suppl. Table S1). Other gene expression platforms like Oncomine and Cosmic were also consulted for additional analysis.

### Seahorse Extracellular Flux Analysis

Mitochondrial respiration in SK-MEL-28, SK-MEL-2, SK-MEL-5, UACC-257, UACC-62 and RPE cells was characterized by extracellular flux analysis using Agilent Seahorse XF96 Analyzer (Agilent Seahorse Bioscience, Santa Clara, CA, USA), as an indicator of the functional bioenergetic capacity of the mitochondria and overall cellular health in response to TRMT61B downregulation.

To this end, the indicated cell lines were subjected to Agilent Seahorse XF Cell Mito Stress Test, and oxygen consumption rate (OCR) was measured in function of time and added ETC disruptors according to the manufacturer’s indications (Ref: https://www.agilent.com/cs/library/usermanuals/public/XF_Cell_Mito_Stress_Test_Kit_User_Guide.pdf). In short, 5-7 days after the last lentiviral infection, control and TRMT61B interfered cells were harvested and seeded in hexaplicates at the optimized cell density (3×10^5^ for SK-MEL-2, SK-MEL-28 and UACC-62; 2.5×10^5^ for SK-MEL-5; 1.5×10^5^ for UACC-257 and RPE) in a Seahorse XF96 Cell Culture Microplate (Agilent Seahorse XFe96 FluxPak: 102416-100) and were allowed to adhere for 24 h in a 37 ⍰C humidified incubator with 5% CO2. The four corners were left only with medium for background correction. On the day of the analysis, the culture medium was replaced with 175 μl assay medium (bicarbonate-free DMEM supplemented with 2mM Gln, 1mM Pyruvate and 10 mM Glucose), and cells were incubated at 37°C for 60 min before measurement in a non-CO2 incubator. OCR was determined at basal conditions and after addition of different ETC disruptors: oligomycin (1 μM) was used to block ATP synthase, carbonyl-cyanide-4-(trifluoromethoxy) phenyhydrazone (FCCP, 0.25 μM) was used to make the inner mitochondrial membrane permeable for protons and allow maximum electron flux through the electron transport chain, and a mix of rotenone (0.5 μM) and antimycin A (0.5 μM) were used together to inhibit complexes I and III, respectively. Basal respiration is determined as the oxygen consumption rate used at basal conditions in the presence of glucose and pyruvate. ATP-linked respiration is the difference between the basal respiration and the resulting OCR after inhibition of the ATP synthase by oligomycin. Maximal respiration is the maximal OCR attained by addition of the uncoupler FCCP and shows the maximum rate of respiration that the cell can achieve. Spare respiratory capacity indicates the capability of the cell to respond to an energetic demand and is the difference between the maximal and the basal respiration. The results shown in Figure 5 are derived from 5 independent experiments.

For the analysis of the glycolytic capacity of the cells XF Glycolysis stress kit was used (Ref: https://www.agilent.com/cs/library/usermanuals/public/XF_Glycolysis_Stress_Test_Kit_User_Guide.pdf). Briefly, cells were seeded and handled as previously described. For the assay, growth media was replaced by minimal media (bicarbonate-free DMEM supplemented with 2mM Gln) 1h before the analysis. Glucose (10 mM) was injected, followed by oligomycin (1 μM), which as an ATP inhibitor permits the readout of the maximal ECAR (extracellular acidification rate). Finally, 2-DG (50 mM) was added to completely block glycolysis. The level of glycolysis is calculated as the ECAR rate reached after injection of a saturating amount of glucose. The glycolytic capacity is the maximum ECAR rate reached after shutting down oxidative phosphorylation by oligomycin. The glycolytic reserve is calculated as the difference between the glycolytic capacity and the glycolysis. Data represented in Suppl. Fig. S5 come from 5 independent experiments. Reagent details are included in Suppl. Table S2.

### Analysis of mitochondrial membrane potential

4000-5000 cells lacking or not TRMT61B were seeded in triplicates in 96 well plates. 24 hours later, cells were incubated at 37ºC for 30’ with 50 nM teramethylrhodamine methyl ester (TMRM), a cationic red-orange fluorescence dye that accumulates in active mitochondria with intact membrane potential, in combination with DAPI (10μg/ml) and Mitotracker Green (50 nM) for nuclear staining and mitochondrial mass evaluation, respectively. At least 8 different images per cell line and condition were acquired using a SP5 confocal microscope. The 560 nm laser line was used to excite the TMRM and the 488nm to excite Mitrotracker Green. Emitted fluorescence was measured above 580 nm for TMRM and between 500 and 540 nm for Mitotracker Green. TMRM and Mitotracker Green quantification (calculated as mean intensity) of 2-3 different fields per well was carried out by Image J software and TMRM/Mitotracker Green ratio was determined. Reagents are listed in Suppl. Table S2.

### Detection of TRMT61B mediated m^1^A in mitochondrial tRNA^leu(UUR)^ and 16S rRNA

For semi-quantitative RT-PCR based detection of m^1^A_58_ and m^1^A_947_ in human mitochondrial tRNA^leu(UUR)^ and 16S rRNA, respectively, 100 ng of small RNA containing samples were annealed with specific reverse primers (tRNA^leu(UUR)^ –R1 and 16S rRNA-R1 for the methylated sensitive PCR reaction or tRNA^leu(UUR)^ –R2 and 16S rRNA-R2 for the control PCR strategy) and retrotranscribed. The resultant cDNA was used as template in semi-quantitative PCR using the primer combinations tRNA^leu(UUR)^ –F1 + tRNA^leu(UUR)^ –R2 or 16S rRNA-F1 + 16S rRNA-R1. Similar to other RNA modifications, the presence of m^1^A_58_ and m^1^A_947_ can partially or totally block reverse transcription. Therefore, considering that R1 primers target the downstream region of the methylated adenosine, cDNAs generated by R1 dependent reverse transcription are less efficiently synthesized and amplified by subsequent PCR reactions using the above primer combinations. In contrast, PCR amplification using tRNA^leu(UUR)^ –F1 + tRNA^leu(UUR)^ –R2 or 16S rRNA-F2 + 16S rRNA-R2 primer pairs is not altered by the presence or absence of m^1^A in cDNAs derived from R2 mediated retrotranscription, since R2 annealing occurs upstream of the methylated adenosine. This last condition was used as reference. Primer sequences are provided in Suppl. Table S2.

### RNA extraction, reverse transcription and semi-quantitative PCR

For total RNA extraction, isolated mitochondrial fractions were lysed in Trizol followed by standard phenol-chloroform extraction protocol. Small and long RNAs were purified using miR Neasy kit (see Suppl. Table S2) following the manufacturer’s instructions. For reverse transcription, total amount of RNA isolated from cells was adjusted to 100 ng/μl in RNase-free water. Then, we mixed 1μl of total RNA with 1μl dNTPs 10mM and 2 μl 20μM reverse primer R1 or reverse primer R2, heat-denatured the mixture at 65°C for 5 min and cooled it rapidly on ice for at least 5 min. With the mixture still on ice, reverse transcriptase was added. Reverse transcription was performed in a total reaction volume of 20 μl at 55ºC for 45 min and then heat inactivated at 70°C for 15 min. We subjected 1μl synthesized cDNA to semiquantitative PCR with the 2X *Green Master Mix* supplemented with DMSO (10%) and 0.5μl of each of the corresponding forward and reverse primers (20μM) to a final volume of 20μl. PCR program includes the following time and temperature parameters: denaturation (Temp: 95°C. Time: 5 min on the initial stage; 30 seconds on rest), annealing (Temp: 50°C for tRNA^leu(UUR)^ –F1 + tRNA^leu(UUR)^ –R2 and 58°C for 16S rRNA-F1/F2 + 16S rRNA-R1/R2. Time: 30 seconds) and extension (Temp: 72°C. Time: 30 seconds; 2min on the last cycle). 26 cycles were performed in all the cases.

### Mitochondrial isolation

Around 20-25 ×10^6^ cells growing in four 15 cm plates were collected in 5.2 ml of fractionation buffer (20mM HEPES (pH 7.4), 10mM KCl, 2mM MgCl_2_, 1mM EDTA, 1mM EGTA and protease inhibitor cocktail). Then, cell suspension was passed through a 27-gauge needle 10 times, left on ice for 20 min and centrifuged at 720g during 5 min to pellet nuclei and cell debris. 90% of the supernatant containing cytoplasm, membranes and mitochondria was transferred (the remaining 10% was discarded to minimize contamination) into a fresh tube and centrifuged at 3000 g during 15 min to separate mitochondria (pellet) from cytoplasm fraction (supernatant). Nuclear and mitochondrial fractions were subjected to an additional wash involving syringe-based homogenization and centrifugation at the previously recommended speeds. Cytoplasm was also centrifuged once again (3000 g during 15 min) and pellet was discarded. Mitochondrial fractions were equally divided for ulterior RNA and protein analysis and preserved at −80°C. WB analysis of different nuclear (LaminA/C), mitochondrial (TOMM40) and cytosolic (a-tubulin) markers confirmed an efficient subcellular fractionation with reduced contamination levels in each of the fractions.

### Cell viability assays

Cells from the different cell lines reported along the present study were seeded at 500-1000 cells per well in 96-well, black-walled, clear-bottomed tissue culture plates (see Suppl. Table S2), incubated at 37 °C and monitored every 2-3 days for cell death and proliferation during a minimum period of 7 days. For total and dead nuclei counting assays, Hoechst 33342 and Propidium Iodide were added at 10μg/ml and 1μg/ml final concentration, respectively, and incubated for 15-30 min in the dark at 37 °C. High-throughput quantification was performed by automated imaging system (Cytell, GE Healthcare, Buckinghamshire, UK) equipped with built-in enumeration software and suitable fluorescence filters. Reagents are detailed in Suppl. Table S2.

### Invasion assay

Invasion assays of the indicated melanoma and control cell lines were performed using transwell matrigel coated inserts (see Suppl. Table S2) as previously described in Salvador et al., 2017 with minimal changes ^93^. In brief, 0.7 ml DMEM supplemented with 10% FBS was added to the well of the plate as attractant. Then, proper cell numbers per cell line were seeded by triplicates on the upper part of each chamber in low serum conditions (0.5% FBS). After 48 hours, non-invading cells on the upper surface of the membrane were wiped with a cotton swab and invading cells on the lower surface of the nucleopore membrane were fixed (1% glutaraldehyde), stained with crystal violet (0.5% in PBS) and counted by examination of five microscopic fields per insert. For each cell line, numbers were normalized to the corresponding PLKO-scramble or empty vector (EV) values.

### Senescence analysis

β-Galactosidase activity was measured according to the manufacturer’s instructions (see Table XX). β-Galactosidase cells were determined using a LEICA DM IL LED Inverted Phase Contrast Microscope and the ImageJ 2.0.0 software (NIH, USA) was used for further analysis. 10 different fields were examined per cell line and condition. Percentages of senescence cells were calculated per each cell line and normalized to the corresponding control values. Reagents are listed in Suppl. Table S2.

### Measurement of autophagic flux

Control and TRMT61B deficient SKMEL28, SKMEL2 and SKMEL5 cell lines were treated during 7 hours with the lysosomal protease inhibitor cocktail E64d (5μ/ml) and pepstatin A (5μ/ml) or DMSO vehicle alone. Overall LC3B-II levels were determined by western blot analysis and measured from the scanned film by densitometric analysis using ImageJ software (NIH, USA). The ratio of the LC3B-II protein level to the αTUBULIN was reported as the result of LC3B-II quantification for each experimental condition.

### Expression constructs, shRNAs and Lentiviral transduction

TRMT61B depletion by shRNA was achieved by using the PLKO.1 system. All lentiviral plasmids were commercially obtained from Sigma Aldrich. Lentiviruses were obtained as previously shown in Salvador et al., 2017 with minor changes ^93^. Very briefly, 5 μg PLKO.1 shRNA plasmids were cotransfected with 5μg secondary packaging plasmids into HEK293T cells. Media was harvested at 48 and 72 hours post-transfection and stored at −80°C. Target cells were seeded at 8×10^5^ cells per 10 cm plate and 5 ml of virus with 8μg/ml polybrene was added. Cells were subjected to 2-3 consecutive infections to enhance TRMT61B downregulation. Puromycin was used for selection (2-5 μg/ml). In general, 5-6 days after the last infection round, cells were plated for the different experiments. For generation of the doxycycline inducible lentiviral construction, modified SH2 containing a 7 nucleotide loop (SH2-7NT) was cloned in EZ-Tet-pLKO-puro vector following the protocol described in Frank et al., 2017 ^94^. TRMT61B overexpression construct was obtained from genecopoeia plasmid collection and virus generation was conducted with the third-generation packaging system. Reagents and shRNA sequences are shown in Suppl. Table S2.

### TRMT61B knock-out by CRISPR/Cas9 gene edition

For CRISPR/Cas9 knockout of TRMT61B, SKMEL28 cells were transfected with inducible endonuclease Cas9 construction (gently provided by Dr. Iain Cheeseman) and selected in the presence of G418 (1mg/ml) for 21 days. Then, lentiviral plasmids containing sgRNAs targeting exon 1 of TRMT61B locus were transduced into target cells. 3-4 days after puromycin selection, individual clones were isolated by single cell cloning, expanded and tested for TRMT61B deletion by WB analysis in response to doxycycline treatment. sgRNA sequences and plasmids are listed in Suppl. Table S2.

### Immunoblots

Western blotting was performed using standard methods. In brief, cells were lysed in RIPA buffer (50mM Tris, pH7.4, 150mM NaCl, 1% NP40, 0.1% SDS, 0.5% DOC, protease inhibitors, 30mM NaF, 1mM Na_3_VO_4_ and 0.1mM PMSF). Protein concentration was determined by the BCA assay. Equivalent amounts of protein were separated on precast polyacrylamide midi gels and electroblotted onto nitrocellulose membranes. Dry transfer apparatus was run according to manufacturer’s recommendations (Transfer-Blot Turbo System, BioRad). After blocked with 5% milk, blots were then probed with primary (overnight incubation at 4°C) and the appropriate secondary antibodies (1h at room temperature) diluted in 5% milk. Reactions for protein visualization were detected by enhanced chemiluminescence assay. Antibodies and reagents are listed in Suppl. Table S2.

### Histology and immunohistochemical staining

For histopathological analysis, formalin-fixed paraffin embedded (FFPE) blocks were serially sectioned (3 μm thick) and stained with hematoxylin and eosin (H&E). Additional serial sections were used for immunohistochemical (IHC) studies. Immunohistochemistry was performed on tumor xenografts as well as on commercial (melanoma; see Table S2) and homemade tissue microarrays (CCA) including the number of samples indicated in the corresponding figures. Tissue sections were processed in an Autostainer Link (Dako), with antigen retrieval at pH 9.0 and incubation with TRMT61B, ki67 and cleaved capase-3 primary antibodies (Table S2). Antigens were visualized using 3,3-diaminobenzidine tetrahydrochloride plus (DAB+). A semi-quantitative histology score was calculated considering the intensity of the staining and the number of cells: negative (0), mild (+1), moderate (+2) and strong (+3).

### TRMT61B detection in plasma samples by ELISA

Determination of TRMT61B levels in 1:10 diluted plasma samples from patients with colorectal, breast, lung or ovarian cancer was assessed by ELISA (Suppl. Table S2) following manufacturers’ instructions. Briefly, reagents were brought to room temperature and 100 μl of the standard, control, and plasma samples were added. The plate was then carefully sealed and incubated for 2 h at 37 °C. After washing three times with 350 μl of the wash buffer solution, 100 μl of the detection reagent A was added to each well and incubated for 1 h at 37 °C. After another three washing steps, 100 μl of the detection reagent B were added to each well and incubated for 1 h at 37 °C. Then, wells were washed five and signal developed at 37 °C with 90 μl of TMB substrate per well. Reaction was stopped with 50 μl of stop solution per well and measured at 450 nm on The Spark multimode microplate reader (Tecan Trading AG, Switzerland).

### Xenograft experiments

SK-MEL-103 (2,5–3 × 10^6^) and SK-MEL-28 cells (2,5–3 × 10^6^) in 100 μL of PBS were subcutaneously injected in immunocompromised nod/scid or nude mice, respectively. Tumor dimensions were measured using a caliper and tumor volume was calculated according to the formula V= 4/3πR^2^r. Doxycycline was administrated every 2 days in the drinking water (1mg/ml) with 3% of sucrose from the very beginning (SK-MEL-28 injected cells) or when tumor size reaches 0.15-0.2 cm^3^ (SK-MEL-103 injected cells).

Zebrafish embryos were obtained by mating adult zebrafish (Danio rerio, wild type), maintained in aquaria with a ratio of 1 fish per liter of water, with 14⍰h/10⍰h light/dark cycle and a temperature of 28.5⍰°C according to published procedures ^95^. 10^6^ 10-day doxycycline pretreated or untreated SK-MEL-28 cells harboring iCas9 alone or in combination with sgRNA were resuspended in 10 μl of PBS with 2% of polyvinylpyrrolidone (to avoid aggregation) and injected into the circulation of 48 hpf embryos (100-200 cells per embryo) using a microinjector (IM-31 Electric Microinjector, Narishige) with an output pressure of 34⍰kPA and 30⍰ms of injection time per injection. Inoculated embryos were incubated at a temperature of 34°C in the presence or absence of doxycycline (1μg/ml) and images were captured using a fluorescence stereomicroscope (AZ-100, Nikon) at 1, 4, and 6 dpi (only 1 and 6 days are shown) to measure dissemination and proliferation of the injected cells in the caudal hematopoietic tissue (CHT) of the zebrafish embryos in each of the conditions assayed. The image analysis of the injected embryos was carried out using the Quantifish software v2.1 (University College London, London, UK) to obtain the proliferation ratio of the cells in the region of the caudal hematopoietic tissue of the embryos, where the cells proliferate and metastasize. This program measures in each of the images provided the intensity of the fluorescence and the area of the positive pixel above a certain threshold of the cells. With these parameters, an integrated density value is obtained allowing the researcher to compare different times between images to reach a proliferation ratio. A ratio above and below 1 means cell proliferation and cell death, respectively. Apart from that, the number of tumors present in each of the images corresponding to the tails of the embryos was calculated. Tumor counting was performed based on tumor size and separation inside the tail of the fish.

### Statistics

Statistical analyses *p*-values were generated using Student’s *t*-test (unpaired, 2-tailed). In all figures, \**P* ≤ 0.05, \*\**P* ≤ 0.01, \*\*\**P* ≤ 0.001 and ns *P* > 0.05 p-value. Error bars were calculated as standard error unless otherwise indicated. Sample size and number of experiments are indicated in each of the Figures.

## Supplementary information

**Figure S1.**
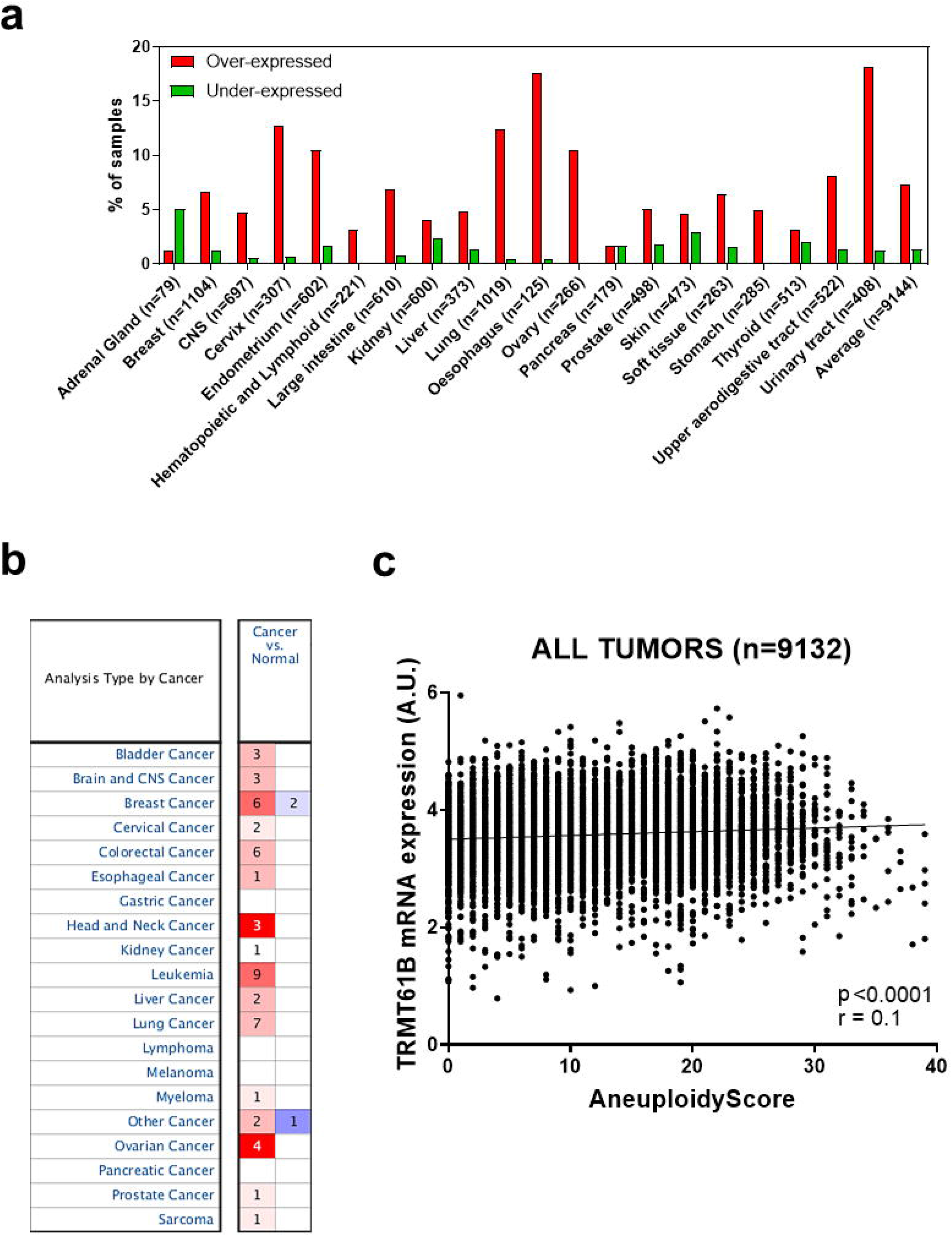
TRMT61B mRNA overexpression in human cancers. TRMT61B is found overexpressed in human cancers according to different bioinformatic platforms. **a** Using Cosmic, TRMT61B mRNA expression was found over-expressed in 19 out of 21 tumor types (https://cancer.sanger.ac.uk/cosmic/gene/analysis?ln=TRMT61B). **b** Similar results were obtained in Oncomine (https://www.oncomine.org/resource/main.html#dso%3AgeneOverex%3Bec%3A%5B2%5D%3Bepv%3A150001.151078%2C3508%3Bg%3A55006%3Bpg%3A1%3Bpvf%3A206439%3Bscr%3Asummary%3Bth%3Ag100.0%2Cp1.00E-4%2Cfc0.0%3Bv%3A18), where 16 out of 20 tumor types showed at least one study with TRMT61B over-expression in cancer samples when compared with control ones (threshold P-value=1E-4). **c** Scatter Plot showing significant correlation between TRMT61B mRNA expression and aneuploidy levels in TCGA dataset (9131 tumors from 33 tumor-types) (Person’s r=0.1, p <0.0001).

**Figure S2.**
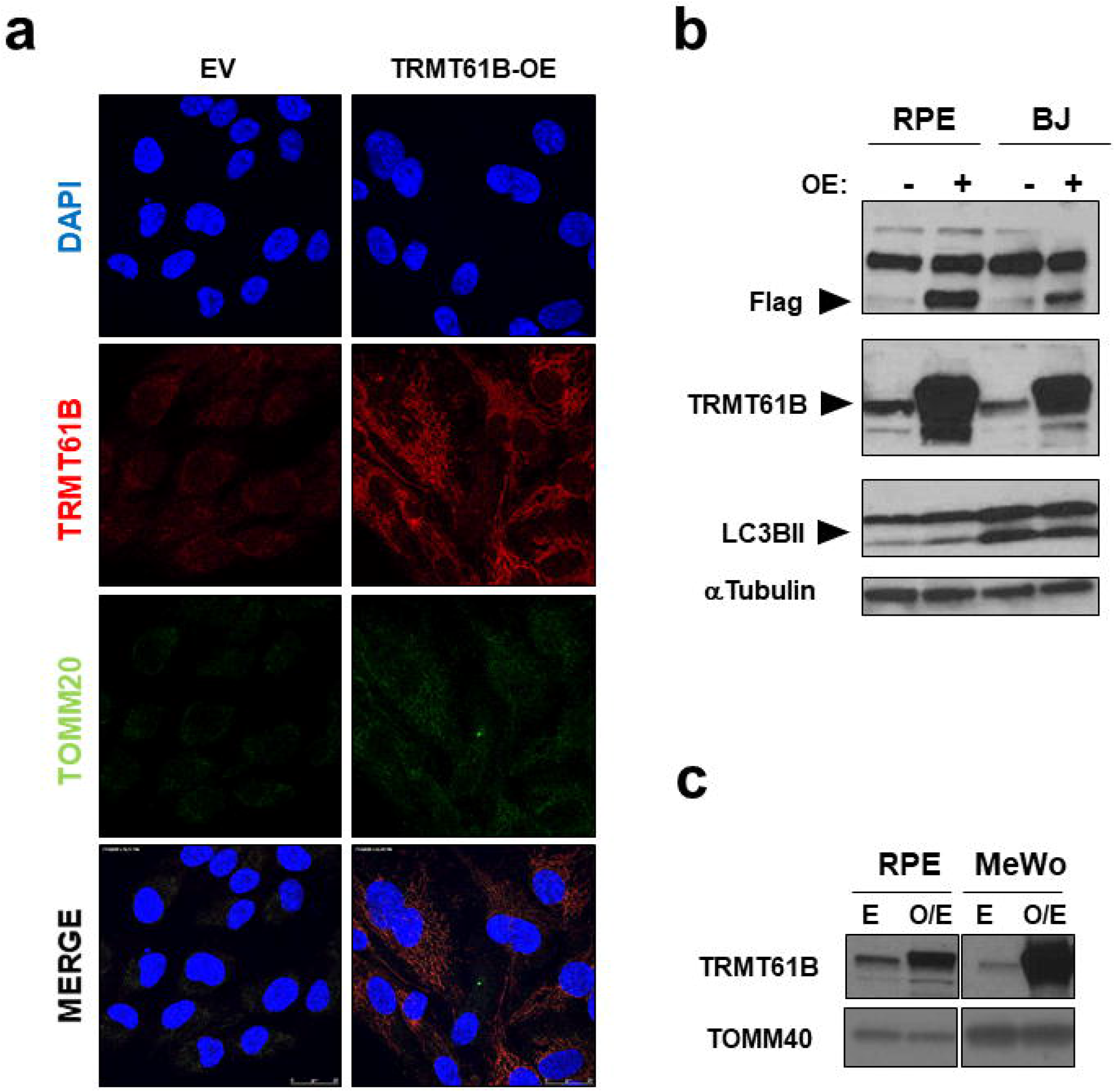
TRMT61B overexpression in euploid and melanoma cell lines. **a** Immunofluorescence analysis of TRMT61B (red) in control and overexpressing RPE cells confirms the previously described mitochondrial localization according to its overlapping expression pattern with the mitochondrial marker TOMM20 (green). Total **(b)** and mitochondrial **(c)** protein extracts obtained from control and TRMT61B overexpressing RPE, BJ and MeWo cell lines were analyzed by western blot for the indicated expression markers.

**Figure S3.**
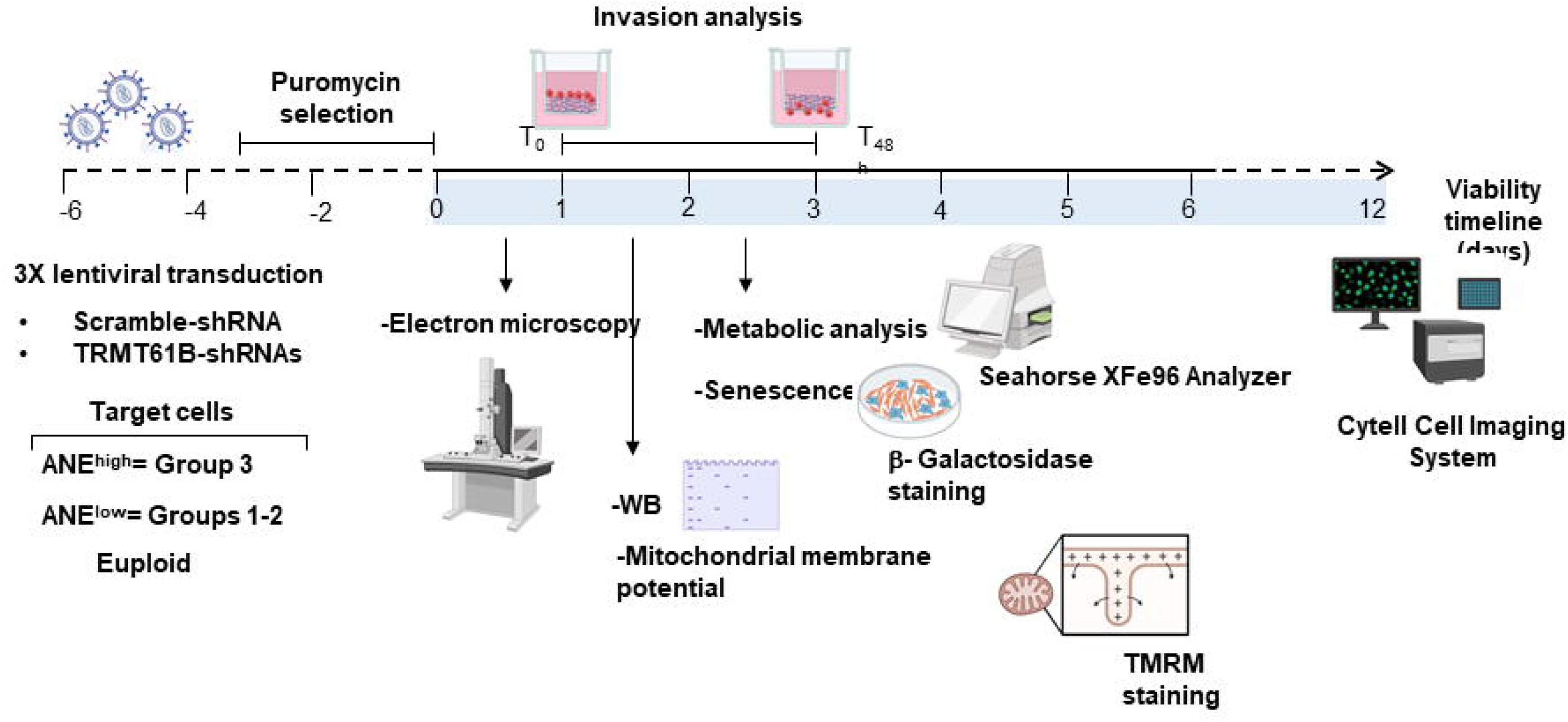
Flow chart detailing the experimental approach designed to evaluate the biological effects upon TRMT61B silencing by shRNA technology. Cells showing different aneuploidy levels were consecutively infected with lentiviral particles harboring scramble or TRMT61B-specific shRNAs. After puromycin selection, cells were subjected to the depicted experimental procedures at the indicated time points.

**Figure S4.**
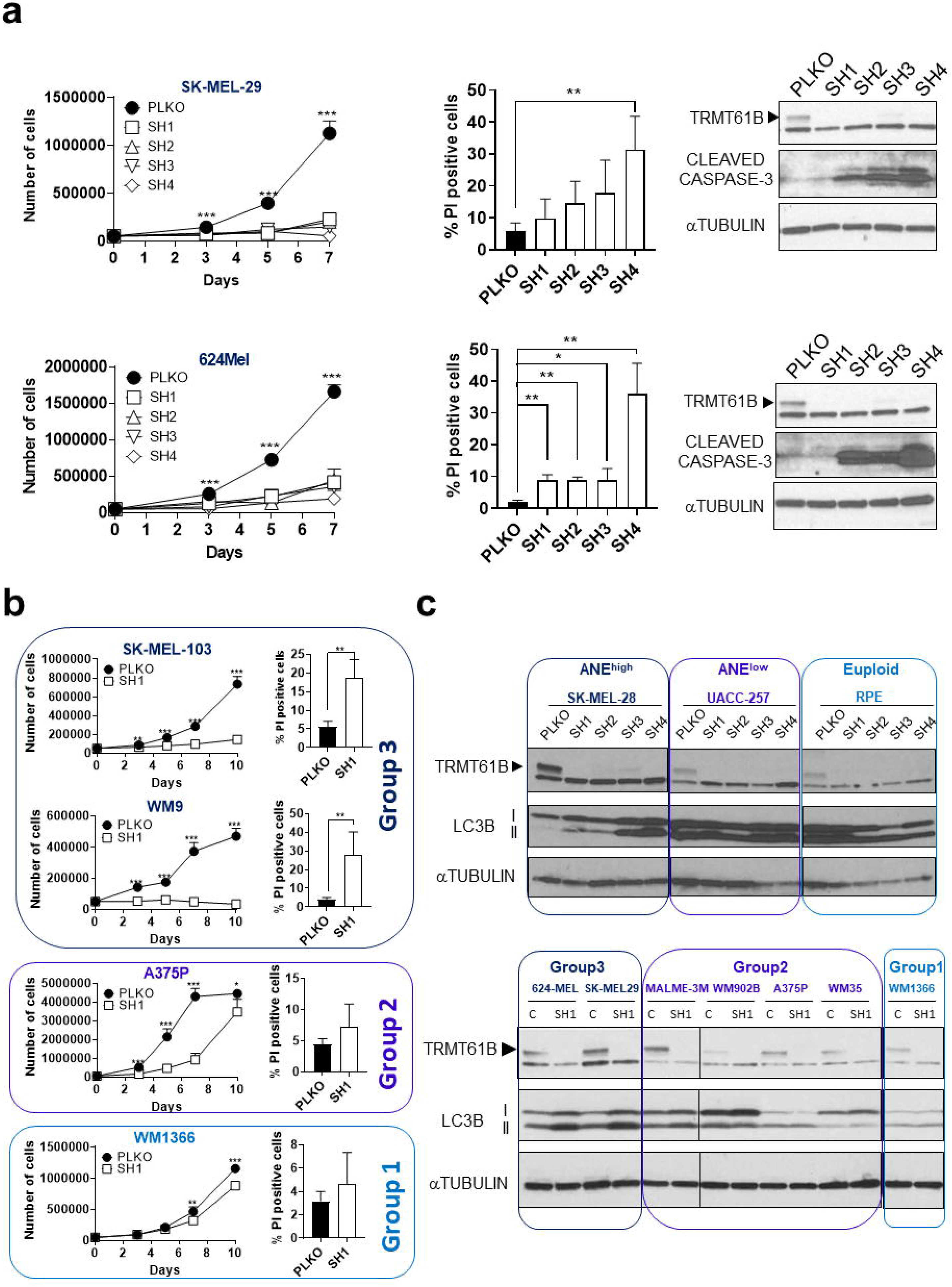
TRMT61B deficiency impacts on proliferation, apoptosis and autophagy more remarkably in ANE^high^ cells. **a** 624-Mel and SK-MEL-29 cell lines exhibit reduced proliferation rates (left graphs) and increased apoptotic levels (middle histograms) in response to TRMT61B elimination. Expression of Cleaved Caspase-3 by western blot analysis is also shown in both genetic contexts (right panels). **b** Reduced proliferative activity and increased apoptotic susceptibility of WM9 and SK-MEL-103 cell lines (Group 3 showing the highest aneuploidy levels) compared with A375P and WM1366 cell lines (Groups 1 and 2 with lower aneuploidy levels) after TRMT61B removal. **c** LC3BII expression was determined by western blot in a collection of melanoma cell lines with different aneuploidy ratios. Four (upper) or one (lower) shRNAs were tested in two different assays. Loss of TRMT61B results in a more pronounced increase of LC3BII levels in cells with the highest aneuploidy ratios. Error bars represent standard deviation in all the panels. \**P*< 0.05; **0.001<*P*< 0.01; *** *P*< 0.001 (Student’s *t*-test; unpaired, 2-tailed).

**Figure S5.**
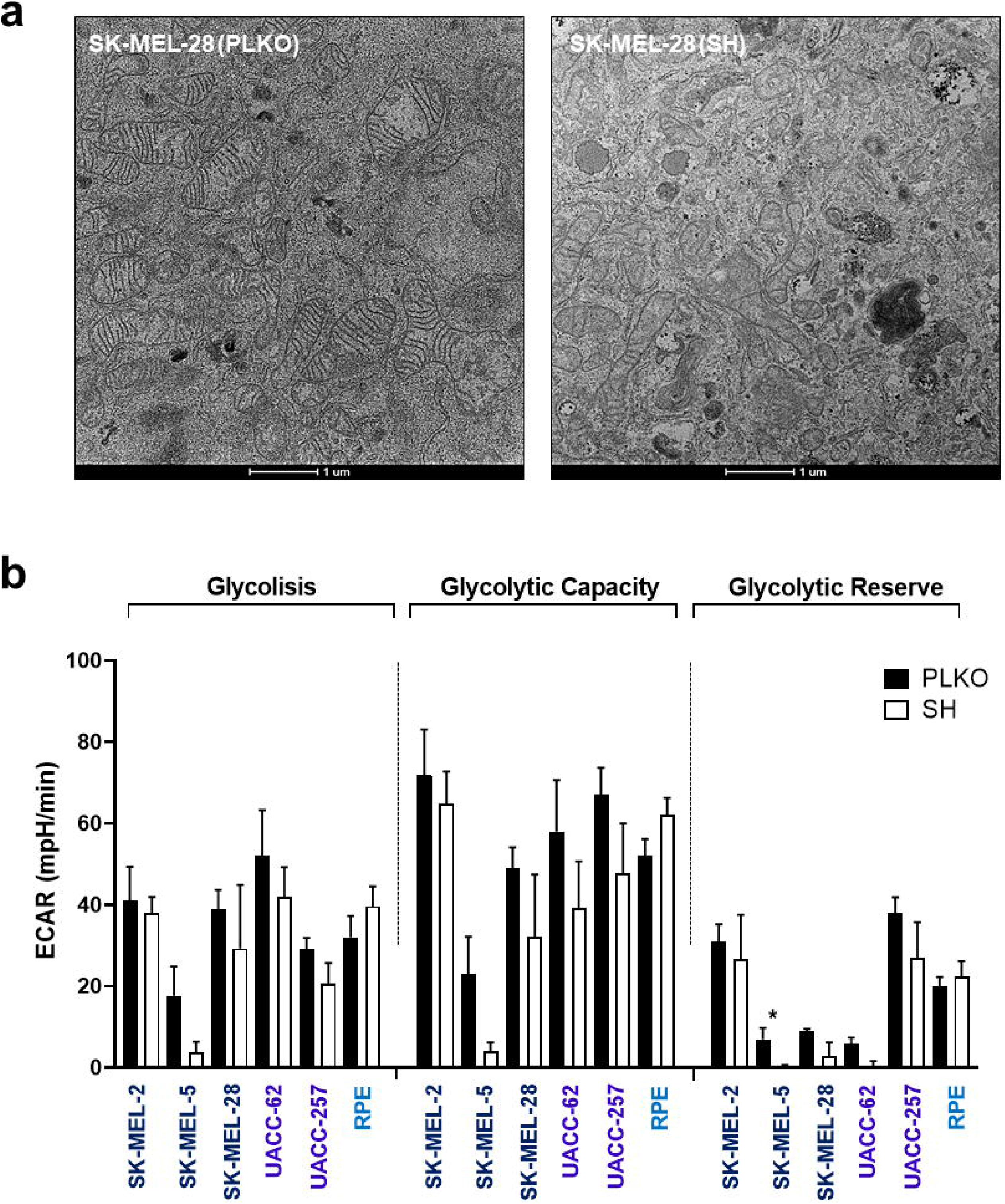
Structural mitochondrial alterations and absence of glycolytic compensation upon TRMT61B depletion. **a** Representative transmission electron micrograph of SK-MEL-28 control and TRMT61B deficient cells showing a high proportion of mitochondria with reduced crest content in the latter. **b** Seahorse analysis of glycolysis in ANE^high^, ANE^low^ and euploid cells does not reflect significant changes when TRMT61B is depleted. Continuous ECAR values (mpH/min) are shown. Glycolytic functions were estimated as explained in Supplemental material and methods. Five independent experiments were carried out between five and seven days after TRMT61B silencing using two different shRNAs (SH1 and SH4). Error bars represent standard error. *p⍰<⍰0.05.

**Figure S6.**
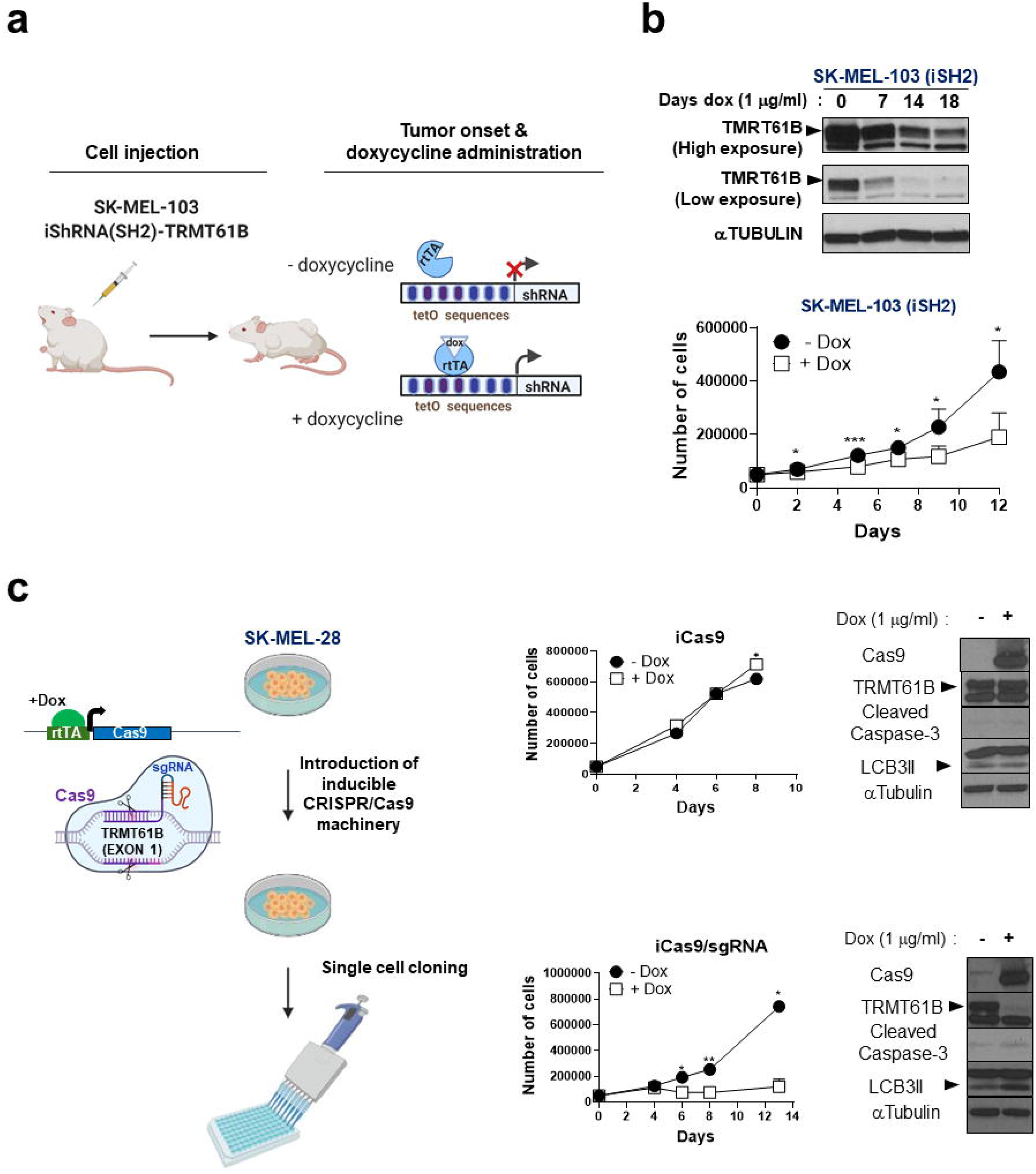
Impairment of cell proliferation after inducible disruption of TRMT61B. **a** Therapeutic strategy to analyze antitumoral activity mediated by TRMT61B suppression in established tumor lesions is represented in the scheme. SK-MEL-103 cells were infected with a doxycycline inducible shRNA construction (tet-on system), subjected to puromycin selection and finally subcutaneously injected in NOD-SCID mice. When tumors reach 0.150-0.200 cm^3^ doxycycline is administrated in the drinking water to the mice. **b** SK-MEL-103 cells were transduced with an inducible shRNA construction (iSH2) specific for TRMT61B and treated with doxycycline during several days. Protein extracts from treated and non-treated cells were examined for TRMT61B expression at different time points showing a dramatic reduction in the presence of doxycycline (left panel). 7 days after treatment, cell proliferation started to be evaluated by Cytell analysis with a negative impact in response to doxycycline administration (right graph). Error bars represent standard deviation. \**P*< 0.05; **0.001<*P*< 0.01; *** *P*< 0.001 (Student’s *t*-test; unpaired, 1-tailed). One representative experiment is depicted. **c** Doxycycline inducible Crispr/Cas9 machinery was sequentially introduced in SK-MEL-28 cells by transfection (inducible Cas9) and infection (specific sgRNAs or empty vector). Cell cultures harboring endonuclease Cas9 alone (iCas9) or both CRISPR elements (iCas9/sgRNA) were subjected to single cell cloning to select proper clones (left diagram). TRMT61B expression is efficiently abrogated only when both Cas9 and TRMT61B specific sgRNAs coexist in the system (iCas9/sgRNA +Dox) inducing a profound proliferation decay. In sharp contrast, untreated iCas9/sgRNA and iCas9 cells regardless of doxycycline regimen, proliferate with normal kinetics. Western blot analysis of relevant markers is also shown. Two independent experiments were performed. Error bars represent standard error. \**P*< 0.05; **0.001<*P*< 0.01; *** *P*< 0.001 (Student’s *t*-test; unpaired, 2-tailed). One representative experiment is depicted.

**Figure S7.**
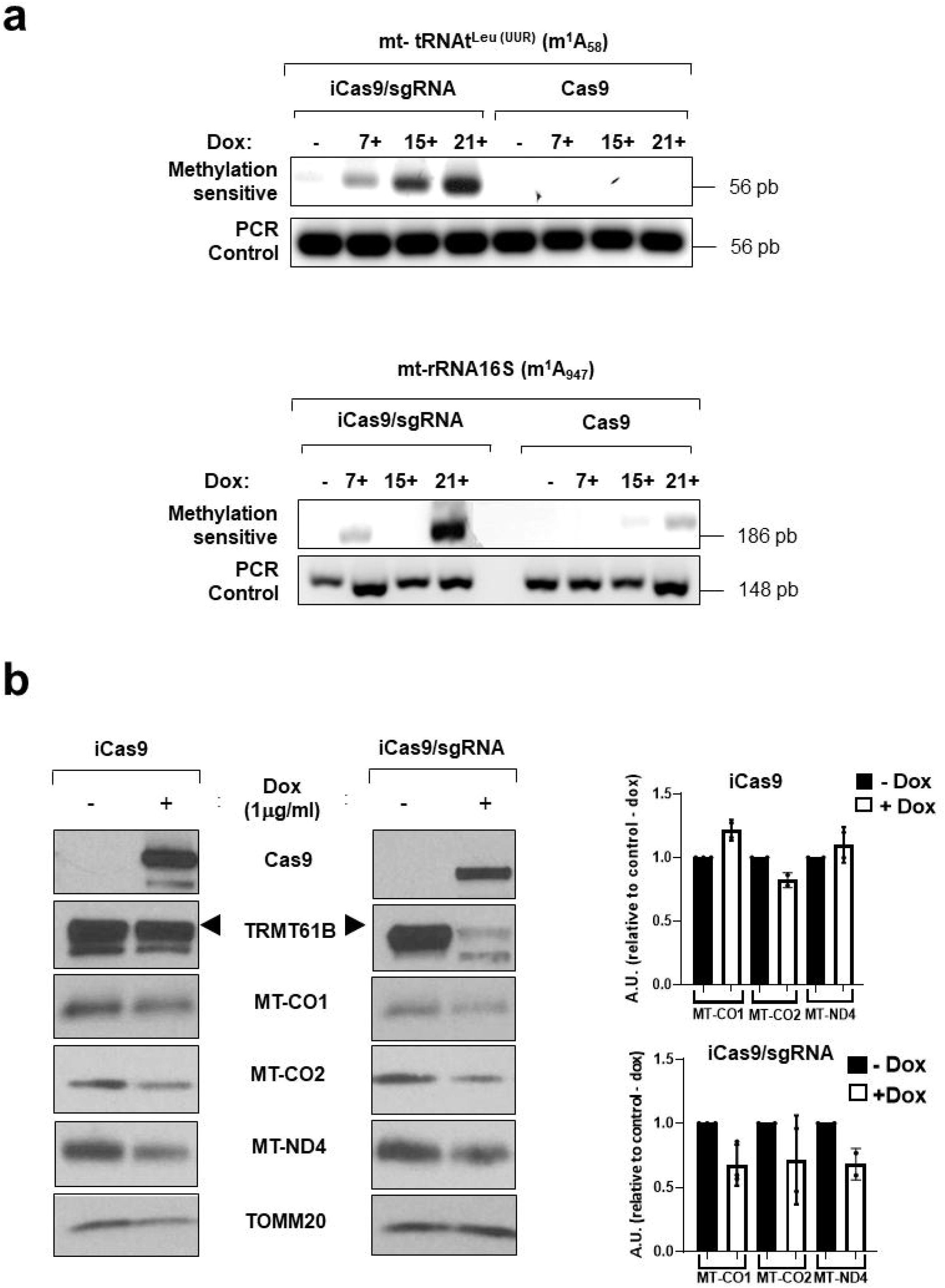
Crispr/Cas9 mediated loss of TRMT61B decreases specific m^1^A methylation and expression of several mitochondrial encoded genes. **a** Two well-known TRMT61B mt-RNA targets were assayed for their methylation levels in the indicated adenosine residues (see brackets) following the PCR strategy described in Material and Methods. TRMT61B removal in doxycycline treated iCas9/sgRNA cells leads to a significant reduction in the amount of methylated RNAs according to the increased cDNA amplification detected along the time (7, 15 and 21 days post-doxycycline treatment). **b** Representative western blot analysis (left panels) of different mitochondrial protein coding genes is shown. Quantification of 2-3 independent experiments (right histograms) indicates a gene expression decrease when TRMT61B is eliminated.

## Acknowledgments

We acknowledge master thesis’ students Martín Salamini and Sarai Araujo for their support in the performance of several *in vitro* experiments. We thank research groups led by Amparo Cano, Marisol Soengas and Marcos Malumbres for providing us most of the cell lines used in this study. We would also like to thank Daniel Luque and María del Carmen Terrón, members of the Electron Microscopy Unit from ISCIII, for their technical assistance. All the illustrations included in the figures were created with the help of biorender.com. A.M. is a recipient of a postdoctoral fellowship from the Fundación Española Contra el Cáncer (AECC). This study was supported by grants from the Spanish Ministry of Economy and Innovation (to I.P.C., SAF2016-76929-R) and from the Acción Estratégica de Salud Intramural (to A.M., PI17CIII/00010).

## Author contributions

A.M. acquired, analyzed and interpreted most of the data; B.-V., I.H., D.G.-D and S.A. assist with several *in vitro* experiments; B.-V. and C.E. helped with mouse xenograft experiments; A.C. and S.V. carried out the Seahorse studies; R.IR.M provided liver samples and performed the subsequent histological analysis; E.C. is our in-house pathologist and assisted with histological analysis of melanoma TMA and xenograft tumors; R.B. and M.G.A. performed ELISA assays and collected serum from normal and colon patients; J.C. provided additional serum samples from breast, ovary and lung tumor bearing patients; A.M.-R. performed karyotype analysis; P.C.-S. and L.S designed and performed the zebrafish studies. I.P.C and A.M. developed the study concept, obtained funding, interpreted the data and drafted/edited the manuscript; and all the authors edited the manuscript.

