## Supplementary material for "Targeting the mitochondrial RNA methyltransferase TRMT61B reveals new therapeutic opportunities in aneuploid cancer cells": Table S2

**Supplementary Table 2.** Key resources used in this work.

| **ANTIBODIES** | | | | | | |  |
| --- | --- | --- | --- | --- | --- | --- | --- |
| **Name** | | | | **Source (identifier)** | **WB** | **IF** | **IHC** |
| Anti-rabbit TRMT61B | | | | Sigma-Aldrich (St. Louis, MI, USA) (HPA026751) | 1:500 | 1:500 | 1:200 |
| Anti-mouse α-tubulin | | | | Sigma-Aldrich (St. Louis, MI, USA) (T9026) | 1:2500 |  |  |
| Anti-mouse DDDDK (equivalent to FLAG antibody) | | | | Abcam (Cambridge, UK) (ab49763) | 1:1000 |  |  |
| Anti-rabbit LC3B | | | | Cell Signaling (Danvers, MA, USA) (#2775) | 1:1000 |  |  |
| Anti-rabbit Cleaved Caspase-3 (5A1E) | | | | Cell Signaling (Danvers, MA, USA) (#9664S) | 1:500 |  |  |
| Anti-rabbit p21 | | | |  | 1:500 |  |  |
| Anti- rabbit MTCO2 (EPR3313) | | | | Abcam (Cambridge, UK) (ab109739) | 1:400 |  |  |
| Anti- mouse MTCO1 (5D11-1C9) | | | | Abcam (Cambridge, UK) (ab219824) | 1:400 |  |  |
| Anti-mouse MT-CYB (5B3-6E3) | | | | Abcam (Cambridge, UK) (ab219823) | 1:1000 |  |  |
| Anti-rabbit MT-ND1 | | | | Abcam (Cambridge, UK) (ab222892) | 1:2000 |  |  |
| Anti-mouse ND4 (9E4-2D8) | | | | Abcam (Cambridge, UK) (ab219822) | 1:400 |  |  |
| Anti-rabbit ND6 | | | | Abcam (Cambridge, UK) (ab81212) | 1.400 |  |  |
| Anti-rabbit TOMM40 (EPR6932) | | | | Abcam (Cambridge, UK) (ab185543) | 1:1000 |  |  |
| Anti-mouse TOMM20 | | | | Abcam (Cambridge, UK) (ab56783) | 1:500 | 1:300 |  |
| Anti-rabbit MRPL11 | | | | Proteintech (Rosemont, IL, USA) (15543-1-AP) | 1:1000 |  |  |
| Anti-rabbit MRPS27 | | | | Proteintech (Rosemont, IL, USA) (17280-1-AP) | 1:1000 |  |  |
| Anti-Ki67 | | | | DAKO (Santa Clara, USA) (IR626) |  |  | 1:200 |
| Anti-Cleaved Caspase-3 (asp175) | | | | Cell Signaling (Danvers, MA, USA) (#9661) |  |  | 1:200 |
| Anti- mouse CRISPR-Cas9 (7A9-3A3) | | | |  | 1:2000 |  |  |
| anti-Rabbit IgG, HRP-linked whole antibody (from donkey) Secondary Antibody | | | | GE Healthcare (Chicago, Illinois, USA) (NA934V) | 1:2500 |  |  |
| anti-Mouse IgG, HRP-linked whole antibody (from sheep) Secondary Antibody | | | | GE Healthcare (Chicago, Illinois, USA) (NA931V) | 1:2500 |  |  |
| Goat anti-rabbit Alexa Fluor 594 | | | | Thermo Fisher Scientific (Waltham, MA, USA) (A32740) |  | 1:500 |  |
| **CELL CULTURE MEDIA** | | | | **REAGENTS and PLASMIDS (cont.)** | | |  |
| DMEM (Dulbecco’s modified Eagle’s medium) high glucose / Invitrogen (Waltham, MA, USA) (#61965-026) | | | | LV203 containing TRMT61B cDNA/ Genecopoeia (Rockville, MD,USA) (EX-A3710-LV203) | | |  |
| FBS (fetal bovine serum) Tetracycline Negative / Capricorn Scientific (Ebsdorfergrund Germany) (FBS-TET-12A) | | | | LV203 empty vector / Genecopoeia (Rockville, MD,USA)( EX-NEG-LV203) | | |  |
| Doxycycline hyclate/ Formedium (Hunstanton, England) (Dox100) | | | | pVsVg, PLP1 and PLP2 (third generation lentiviral packaging vectors)/Kindly provided by Dr. Amparo Cano | | |  |
| Bicarbonate-free DMEM/ Sigma-Aldrich (St. Louis, MI, USA) (D5030-10L) | | | | pAX2 and pMD2G (second generation lentiviral packaging vectors)/ Kindly provided by Dr. Amparo Cano | | |  |
| FBS (fetal bovine serum) / Sigma-Aldrich (St. Louis, MI, USA) (#F7524-500ML) | | | | SuperScript III First Strand/ Invitrogen (Waltham, MA, USA) (18080-051) | | |  |
| Penicillin/streptomycin / Lonza (Basel, Switzerland) (#DE17-602E) | | | | Lipotransfectin/ Solmeglas (Pozuelo de Alarcón, Madrid, Spain) (SBM 0959) | | |  |
| **SOFTWARE AND PLATFORMS** | | | | ProLong Gold antifade reagent/ Life Technologies (Carlsbad, California, USA) (P36931) | | |  |
| ImageJ (U.S. National Institutes of Health, Bethesda, Maryland, USA) | | | | Puromycin / InvivoGen (San Diego, CA, USA) (ant-pr-1) | | |  |
| Prism 8 (GraphPad Software, Inc) | | | | G418/ InvivoGen (San Diego, CA, USA) (ant-gn-5) | | |  |
| **REAGENTS and PLASMIDS** | | | | DNA polymerase / NZYTech (Lisbon, Portugal) (MB354) | | |  |
| pLenti-sgRNA / Addgene (Watertown, MA, USA) (#71409) | | | | Image-iT TMRM Reagent/ Invitrogen (Waltham, MA, USA) (I34361) | | |  |
| EZ-Tet-PLKO-puro / Addgene (Watertown, MA, USA) (#85966) | | | | MitoTracker Green FM/ Invitrogen (Waltham, MA, USA) (M7514) | | |  |
| Hp-138 (piggybac plasmid for doxycycline dependent Cas9 expression)/ Kindly provided by Dr. Iain Chesseman | | | | BCA system / Pierce (Waltham, MA, USA) (23227) | | |  |
| Transposase expression vector for piggybac insertion/Kindly provided by Dr. Bon-Kyoung Koo’s lab | | | | ECL western blotting system / Thermo Fisher Scientific (Waltham, MA, USA) (Pierce 32106) | | |  |
|  | | | **REAGENTS and PLASMIDS (cont.)** | | | |  |
|  | Pepstatin A/ Sigma-Aldrich (St. Louis, MI, USA) (P5318) | | | | | |  |
|  | E-64d / Sigma-Aldrich (St. Louis, MI, USA) (E8640) | | | | | |  |
|  | TBS (Tris-buffered saline) / Canvax Biotech (Cordoba, Spain) (BR0042) | | | | | |  |
|  | Hoechst 33324 / Thermo Fisher Scientific (Waltham, MA, USA) (H3570) | | | | | |  |
|  | Propidium iodide (PI)/ Sigma-Aldrich (St. Louis, MI, USA) (P5318) | | | | | |  |
|  | DAPI/ Sigma-Aldrich (St. Louis, MI, USA) (D9542) | | | | | |  |
|  | Polybrene/ Sigma-Aldrich (St. Louis, MI, USA) (H9268) | | | | | |  |
|  | Corning Matrigel Invasion Chamber 24-well plate 8.0 Micron/ Corning (Bedford, MA, USA) (354480)  Senescence β-Galactosidase Staining/ Cell Signalling (Danvers, MA, USA) (#9860) | | | | | |  |
|  | Criterion TGX Stain Free Precast Gels/ Biorad (Hercules, California, USA) (5678084)  Transfer-Blot Turbo Transfer Pack/ Biorad (Hercules, California, USA) (1704159)  miRNeasy Mini Kit/ Qiagen (Hilden, Germany) (1038703) | | | | | |  |
|  | Cochicine/ Sigma-Aldrich (St. Louis, MI, USA) (C3915)  Oligomycin A/ Sigma-Aldrich (St. Louis, MI, USA) (75351-5MG)  2-Deoxy-D-Glucose/ Sigma-Aldrich (St. Louis, MI, USA) (D8375-5G)  FCCP / Sigma-Aldrich (St. Louis, MI, USA) (C2920-50MG)  Antimycin A/ Sigma-Aldrich (St. Louis, MI, USA) (A8674-100MG)  Rotenone/ Sigma-Aldrich (St. Louis, MI, USA) (R8875-1G)  96-well plates/ Greiner bio-one (Les Ulis, France) (655986)  Human tRNA Methyltransferase 61B (TRMT61B) ELISA Kit / abbexa (Cambridge, UK) (abx383954)  Human Malignant Melanoma Tissue Microarray / Biomax (Derwood, MD, USA) (ME2082c) | | | | | |  |

**Table 2.** DNA sequences used in this work.

**Name and 5’ to 3’ Sequence**

**sgRNA.1**: TTCGACCTCGGTAGCGGACT

**sgRNA.2**: AGTCCCGTTCGGCAAGATCG

**SH1** (TRCN0000157102): CCGGGCGAGGTCATTGTCAGAGATTCTCGAGAATCTCTGACAATGACCTCGCTTTTTTG

**SH2 (**TRCN0000157325): CCGGCCAGATACTGAGGAGTTCCTTCTCGAGAAGGAACTCCTCAGTATCTGGTTTTTTG

**SH3** (TRCN0000155894: CCGGCCTGTGAACTTGCTCTTTCATCTCGAGATGAAAGAGCAAGTTCACAGGTTTTTTG

**SH4** (TRCN0000151548): CCGGGTCTTGCAGTTAGTTTGACATCTCGAGATGTCAAACTAACTGCAAGACTTTTTTG

**SCRAMBLE:** CTAGCCCTAAGGTTAAGTCGCCCTCGCTCGAGCGAGGGCGACTTAACCTTAGGTTTTTTG

**SH2 (7NT):** CTAGCCCAGATACTGAGGAGTTCCTTTACTAGTAAGGAACTCCTCAGTATCTGGTTTTTTG

**RNAt-Leu(UUR)-R1**: tggtgttaagaagagg

**RNAt-Leu(UUR)-R2**: acctctgactgtaaag

**RNAt-Leu(UUR)-F1**: gttaagatggcagagc

**RNAr 16S-R1**: gggtcttctcgtcttgctgt

**RNAr 16S-R2**: AATCTGACGCAGGCTTATGC

**RNAr 16S-F1**: CCTGCCCAGTGACACATGTT

**RNAr 16S-F2**: TCAAGCTCAACACCCACTACC
